# RanBP2/Nup358 enhances miRNA activity by sumoylating and stabilizing Argonaute 1

**DOI:** 10.1101/555896

**Authors:** Qingtang Shen, Yifan E. Wang, Mathew Truong, Kohila Mahadevan, Jing Ze Wu, Harrison W. Smith, Craig Smibert, Alexander F. Palazzo

## Abstract

RanBP2/Nup358 is one of the main components of the cytoplasmic filaments of the nuclear pore complex. It has been speculated that RanBP2, which has an E3 SUMO-ligase domain, may alter the composition of messenger ribonucleoprotein (mRNP) complexes as they emerge from the nuclear pore and thus regulate the ultimate fate of the mRNA in the cytoplasm. Four separate missense mutations in RanBP2 cause Acute Necrotizing Encephalopathy 1 (ANE1), which manifests as a sharp rise in cytokine production after common viral infections such as influenza and parainfluenza. However, how RanBP2 and its ANE1-associated mutations affect cytokine production is not well understood. Here we report that RanBP2 represses the translation of the *interleukin-6 (IL6*) mRNA, which encodes a cytokine that is aberrantly up-regulated in ANE1. Our data indicates that soon after its production, the *IL6* mRNP recruits the RNA-induced silencing complex (RISC) bound to *Let7* miRNA. After this mRNP is exported, RanBP2 sumoylates the RISC-component AGO1, thereby stabilizing it and enforcing mRNA silencing. Collectively, these results support a model whereby RanBP2 promotes an mRNP remodelling event that is critical for the miRNA-mediated suppression of clinically relevant mRNAs, such as *IL6*.

## INTRODUCTION

Ran-binding protein 2 (RanBP2), also known as Nucleoporin 358 KDa (Nup358), is one of the main components of the cytoplasmic filaments of the nuclear pore complex (NPC) (Hoelz et al., 2011). It has a SUMO E3-ligase domain that post-translationally modifies several proteins (Pichler et al., 2002) and has been implicated in regulating mRNA metabolism (Forler et al., 2004; Grünwald and Singer, 2010; Mahadevan et al., 2013). In particular, mRNAs are known to be packaged into messenger ribonucleoprotein (mRNP) complexes which are thought to undergo maturation events where their proteins are exchanged or post-translationally modified. Virtually all mRNPs must cross the nuclear pore prior to their translation, permitting nuclear pore filament proteins to survey the entire transcriptome and modulate its output. It has been speculated that some nuclear pore proteins, especially those present on the nucleoplasmic and cytoplasmic faces of the pore, regulate mRNP maturation events (Lund and Guthrie, 2005; Palazzo et al., 2013; Saroufim et al., 2015; Smith et al., 2015; Palazzo and Truong, 2016; Fernandez-Martinez et al., 2016), however this is poorly understood.

Previously, we found that RanBP2 was required for the efficient translation of mRNAs that contain signal sequence coding regions (SSCRs), which code for short hydrophobic polypeptides and are found at the 5′ end of the open reading frame (ORF) of most secretory and membrane-bound proteins (Mahadevan et al., 2013). The majority of SSCRs in vertebrates are depleted of adenines, are enriched in GC-motifs and are present in the first exon (Cenik et al., 2011, 2017; Palazzo et al., 2013, 2007). Importantly, human RanBP2 contains eight zinc fingers that directly bind to adenine-depleted SSCRs (Mahadevan et al., 2013). Moreover, the ability of RanBP2 to promote translation is dependent on its zinc fingers and the presence of an adenine-depleted SSCR (Mahadevan et al., 2013). Overall our results suggest that upon the completion of nuclear export, mRNAs that contain adenine-depleted SSCRs directly interact with RanBP2 through its zinc fingers, and that this interaction likely modifies proteins that are associated with the mRNP in order to potentiate the translation of these mRNAs (Mahadevan et al., 2013; Palazzo et al., 2013; Palazzo and Truong, 2016).

Mutations in RanBP2 has also been associated with pathology. In particular, four separate missense mutations in the N-terminal region of RanBP2 (T585M, T653I, I656V, and T681C) are genetic risk factors for a pediatric neurological disease called acute-necrotizing encephalopathy (ANE1) (Neilson et al., 2009; Sell et al., 2016). 40% of individuals with one of these dominant mutations secrete excessive amounts of cytokines (known as a “cytokine storm”) in response to influenza infection (Kansagra and Gallentine, 2011; Singh et al., 2015; Tisoncik et al., 2012). Generally, the massive secretion of cytokines include pro-inflammatory cytokines such as IL6, TNFα, IL10, IFNγ, sTNFα receptor, and IL15 (Akiyoshi et al., 2006; Ichiyama et al., 1998, 2003b, 2003a; Ito et al., 1999; Tabarki et al., 2013; Wu et al., 2015) (Table S1). The resulting elevated levels of cytokines infiltrate into the cerebral spinal fluid, causing neuropathology, seizures, coma and a high rate of mortality. Those who survive often suffer from long-term neurological damage. However, how mutations in RanBP2 contribute to the overproduction of ANE1-associated cytokines remains unclear.

The cytokine that has been best documented to be upregulated during ANE1 is IL6 (Table S1). The expression of this cytokine has been the subject of much investigation. One of the key ways in which IL6 is regulated is by the *Let7* miRNA, which recognizes one binding site in the 3′ untranslated region (UTR) of the *IL6* mRNA (Iliopoulos et al., 2009; Schulte et al., 2011). Indeed, many infections are known to modulate the expression of *Let7* miRNA family members and *Let7* in turn modulates the inflammatory response (Brennan et al., 2017; Iliopoulos et al., 2009; Lin et al., 2017; Ma et al., 2012; Makkoch et al., 2016; Mazumder et al., 2013; Schulte et al., 2011). miRNAs, such as *Let7*, associate with the RNA Induced Silencing Complex (RISC) to silence their targets, and the main component of this complex, the Argonaute (AGO) proteins, are regulated by post-translational modifications, such as ubiquitination and sumoylation, which in turn affect their activity and stability (Chinen and Lei, 2017; Derrien and Genschik, 2014; Josa-Prado et al., 2015; Kobayashi et al., 2018; Nayak et al., 2018; Sahin et al., 2014). Interestingly, a recent report indicates that RanBP2 is required for *Let7* mediated gene silencing (Sahoo et al., 2017). These observations suggest that RanBP2 might impact the translation of the *IL6* mRNA by post-translational regulation of AGO proteins.

Here we present evidence that RanBP2 promotes the *Let7*-mediated suppression of IL6 protein production by sumoylating AGO1, which antagonizes AGO1 ubiquitination and thus promotes its stability and its ability to translationally silence the *IL6* mRNA. Furthermore, we observe that RISC associates with *IL6* mRNA in the nucleus, and then likely accompanies the mRNA through the pore. Our data suggests that when this mRNP reaches the cytoplasm, RanBP2 sumoylates AGO1, which stabilizes this complex and promotes *IL6* mRNA silencing. Thus, our work provides one of the few examples of how mRNPs are subjected to a maturation event at the pore, and how these maturation events affect the ultimate fate of the mRNA in the cytoplasm.

## RESULTS

### Most ANE1-associated Cytokine Genes Have SSCRs that Contain Adenines and Have Low 5IMP Scores

Previously, we found that RanBP2 potentiates the translation of mRNAs that contain adenine-depleted SSCRs at the beginning of their ORF (Mahadevan et al., 2013). Indeed these SSCRs tend to have long tracts of adenine-depleted sequence (Palazzo et al., 2007). Moreover, genes that contain adenine-depleted SSCRs tend to lack introns in their 5′UTRs, when compared to other genes in the human genome (Cenik et al., 2011). As a result, there are typically no introns upstream of adenine-depleted SSCRs, thus placing these elements within the first exon at rates that are higher than expected (Cenik et al., 2011). Genes that contain adenine-depleted SSCRs are also associated with a variety of other features in the 5′ end of the ORF (e.g., presence of certain GC-rich motifs, enhanced presence of N1-methyladenosine), which are also associated with a lack of introns in their 5′UTR (Cenik et al., 2017). Previously, we used machine learning to evaluate the 5′ end of ORFs for these features which are summed up into a <u>5</u>′UTR intron <u>m</u>inus <u>p</u>rediction (5IMP) score (Cenik et al., 2017).

Since ANE1-associated cytokines (see Table S1) are produced from mRNAs that contain SSCRs, we decided to evaluate whether they are also depleted of adenines and have high 5IMP scores. Of note, most of these mRNAs are produced from genes that lack introns in their 5′UTRs (Table S1). Despite this, almost all ANE1-associated cytokines had very small adenine-less tracts when compared to other SSCR-containing genes that lacked introns in their 5′UTRs (“5UI-”) from the human genome (Figure 1A). Indeed, the length of their longest adenine-less tract was on par with non-SSCR containing genes and intergenic sequences (Figure 1A), strongly indicating that there is no selection for adenine-depletion in most ANE1-associated cytokine genes. One exception was IL6, which had a relatively long adenine-less tract. When 5IMP scores were evaluated, all ANE1-associated cytokines, including IL6, had low scores (<5) when compared to other SSCR-containing genes that lack 5′UTR introns (“SSCR 5UI-”) where a significant fraction have scores greater than 7 (Figure 1B).

**Figure 1.**
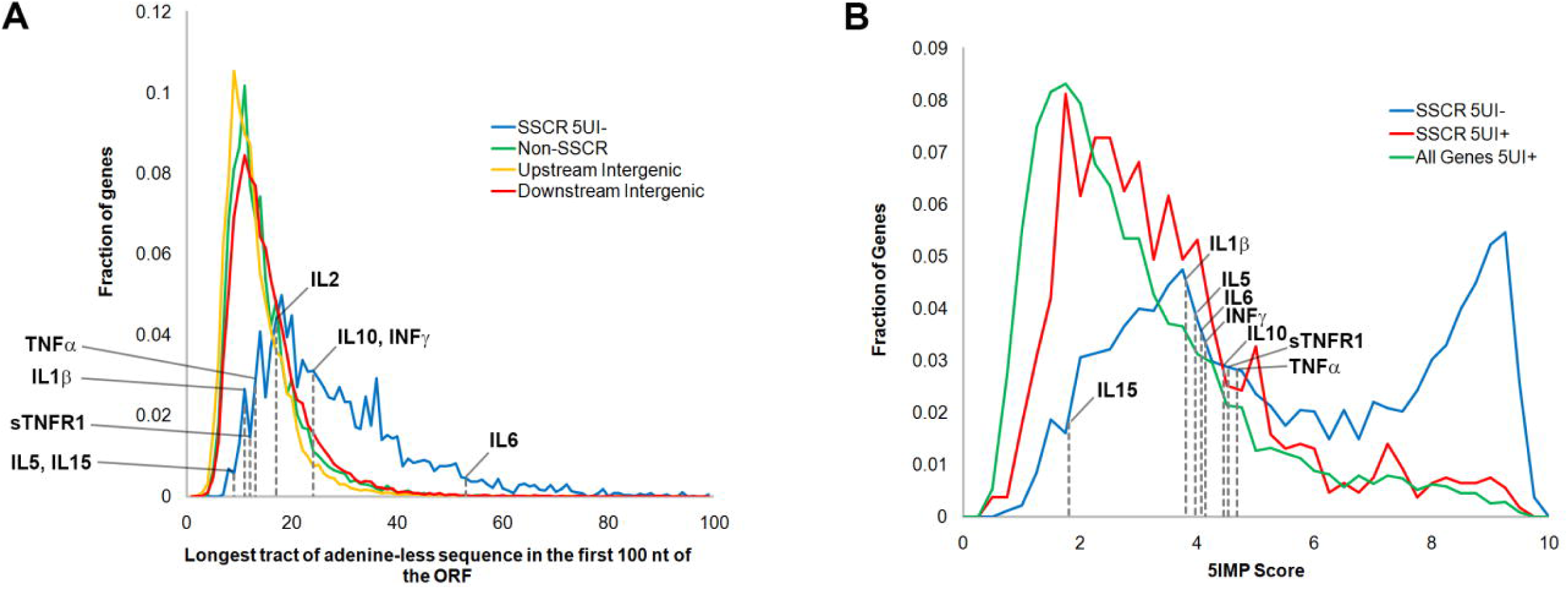
Most ANE1-associated cytokine genes are not depleted of adenines and have low 5IMP scores. (A) For each gene in the human genome, the longest tract of adenine-less sequence in the first 99 nucleotides of the open reading frame was tabulated as in Palazzo *et al,* 2007, and plotted, with the *x-axis* representing the length of these tracts, and the *y-axis* representing the fraction of genes in each set with these tract lengths. This was tabulated for all genes that contain an SSCR that lacks introns in their 5′UTR (“SSCR 5UI-”; blue), which are known to be positively regulated by RanBP2 (Mahadevan et al., 2013), and for genes that lacked SSCRs (“Non-SSCR”; green). To control for the length of adenine-less tracts in random human DNA sequences, the frequency of adenine-less tract length was also tabulated for regions 3 kb upstream (yellow) and 3 kb downstream (red) of protein coding genes. The adenine-less tract lengths for ANE1-associated cytokine genes are labeled. (B) For each gene in the human genome the 5IMP score was calculated, as described in Cenik et al., 2017, and plotted with the *x-axis* representing binned 5IMP scores, and the *y-axis* representing the fraction of genes in each set with these scores. This was tabulated for all genes that contain an SSCR that lacks introns in their 5′ UTR (“SSCR 5UI-”; blue), for genes that contain both an SSCR and one or more introns in their 5′UTR (“SSCR 5UI+”; red) and for all genes that contain one or more introns in their 5′UTR (“All Genes 5UI+”; green). The 5IMP scores for ANE1-associated cytokine genes are labeled.

From these results we conclude that the sequence composition of SSCRs from ANE1-associated cytokine mRNAs do not have the features that are normally associated with mRNAs whose translation is upregulated by RanBP2.

### RanBP2 Inhibits the Translation of an *IL6-HA* Reporter mRNA

Of all the cytokines overproduced in ANE1-patients, IL6 has been the best documented (Table S1). We thus depleted RanBP2 using lentiviral delivered shRNAs (Figure 2A) and examined the expression of a C-terminally tagged IL6 expressed off of a transfected plasmid. Unexpectedly, we found that RanBP2-depletion resulted in a ∼12 fold increase in intracellular IL6-HA when compared to control cells (Figure 2B-C). This was true of both intracellular and secreted IL6-HA (Figure 2D). This was in stark contrast to the expression of the *insulin-HA* reporter, whose mRNA has an adenine-depleted SSCR and a high 5IMP score, and whose translation was reduced in RanBP2-depeleted cells (Figure 2B-C), a result that was consistent with our previous findings (Mahadevan et al., 2013). Protein production from the *βglobin-HA* mRNA, which lacks an SSCR, was unaffected by RanBP2-depletion (Figure 2B-C).

**Figure 2.**
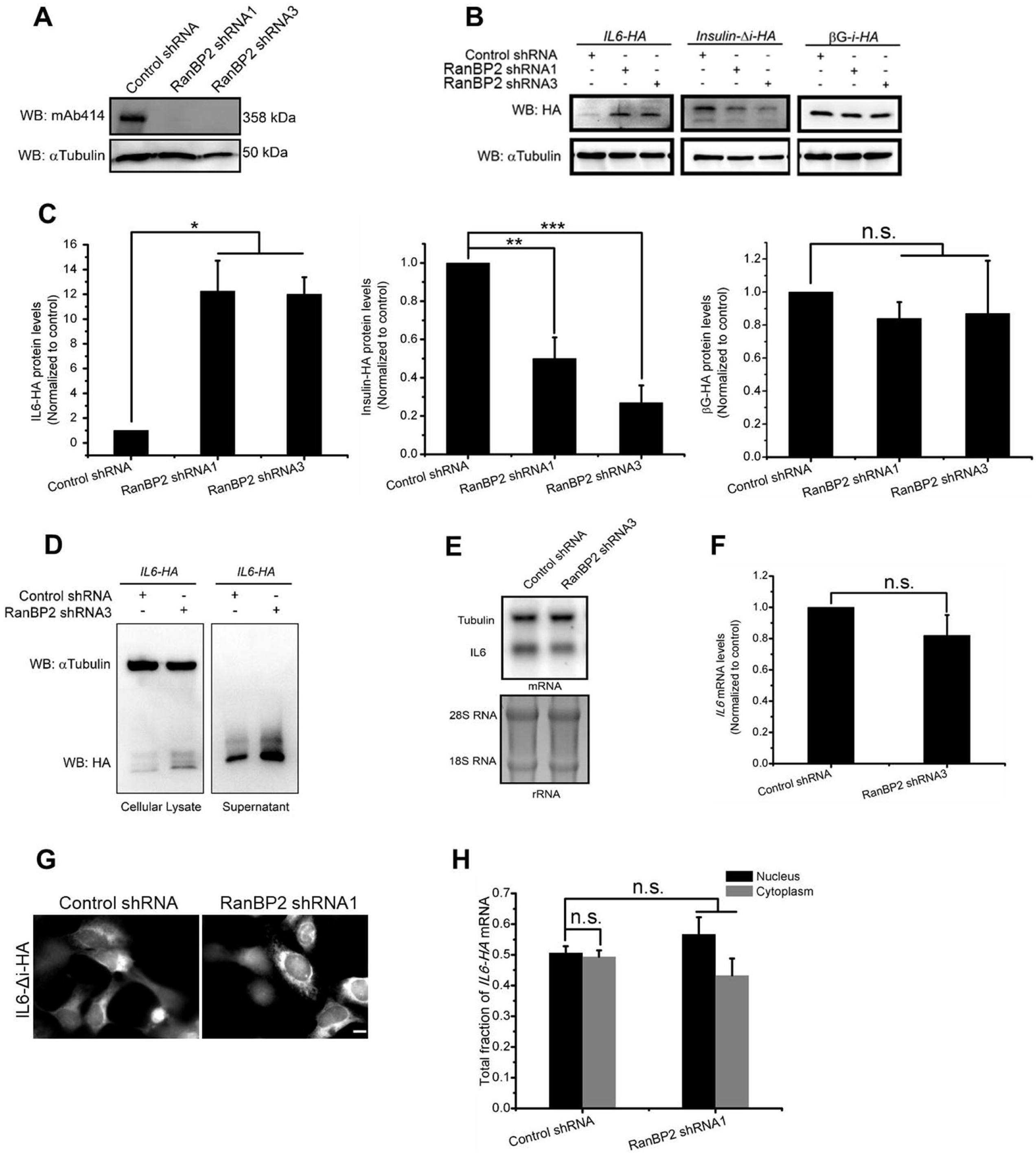
RanBP2 supresses the translation of *IL6* mRNA. (A) U2OS cells were infected with lentivirus containing shRNA1 or shRNA3 directed against RanBP2, or scrambled shRNA (“control shRNA”). Four days post-infection, cell lysates were collected, separated by SDS-PAGE, and immunoblotted for nucleoporins using mAb414, which recognizes RanBP2, and α-tubulin as a loading control. (B-C) U2OS cells were infected with lentivirus that delivered shRNA1 or shRNA3 against RanBP2 or control virus. Three days post-infection, cells were transfected with plasmids containing either the *IL6-HA*, *insulin-HA*, or *β-globin-HA* genes. 18–24 h post-transfection cell lysates were collected and separated by SDS-PAGE. The level of each protein was analyzed by immunoblot for HA, and α-tubulin as a loading control (B). The levels of each HA-tagged protein and α-tubulin were quantified using densitometry analysis. The HA/tubulin ratio was normalized to control shRNA-treated cells and plotted, with each bar representing the average and standard error of three independent experiments (C). (D) As in (B) except that cell lysates (left panel) or supernatant precipitated by TCA (right panel) were collected, separated by SDS-PAGE and immunoblotted with antibodies against HA and α-tubulin. (E-F) As in (B) except that RNA was purified from cell lysates and separated on a denaturing agarose gel. The levels of *IL6-HA* mRNA and *α-tubulin* were assessed by northernblot, while the ribosomal RNA was detected by ethidium bromide (E). *IL6-HA* and *α-tubulin* mRNA levels were quantified using densitometry analysis. The *IL6-HA*/*tubulin* ratio was normalized to control shRNA-treated cells and plotted with each bar representing the average and standard error of three independent experiments (F). (G-H) Control and RanBP2-depleted cells were transfected with *IL6-HA* plasmid for 14-18 hr, then fixed, permeabilized, and stained for mRNA using a fluorescent in situ hybridization (FISH) probe directed against *IL6*. The cells were imaged (G) and total integrated fluorescence was assessed in the cytoplasm and nucleus (H). For each experiment at least 20 cells were assessed with each bar represents the average and standard error of three independent experiments. Scale bar = 10 µm. *P = 0.01–0.05, **P = 0.001–0.01, ***P < 0.001, n.s. indicates no significant difference (Student’s *t*-test).

The increase in IL6-HA protein synthesis was not due to changes in the total level *IL6* mRNA (Figure 2E-F), or to changes in the distribution of *IL6* mRNA between the cytoplasm and the nucleus (Figure 2G-H). Previous studies have demonstrated that splicing can potentiate the efficiency of translation (Nott et al., 2004), and the main *IL6* isoform has four introns. However, versions of the IL6 reporter that either contained the endogenous first intron (*IL6-1i*), or the intron of *fushi tarazu* mRNA (*ftz*), at the first intron site (*IL6-1f*) still produced more protein after RanBP2-depletion, indicating that this effect is independent of splicing (Figure S1A-C).

From these results, and from the results of our polysome profiling (see below), we conclude that RanBP2 inhibits IL6-HA protein production from a transfected reporter construct. As RanBP2-depletion did not affect the levels or the cytoplasmic/nuclear distribution of the reporter mRNA, we concluded that RanBP2 inhibits the translation of *IL6-HA* mRNA.

### The SUMO E3-ligase Domain of RanBP2 Is Required for the Repression of IL6

Next, we investigated whether the SUMO E3-ligase activity of RanBP2 is required to inhibit the translation of the *IL6* mRNA. We used CRISPR/Cas9 with a specific guide RNA (“gRNA-dE3-1”) to target the E3 domain of RanBP2 in U2OS cells (Figure 3A-B). We obtained a clone, called RanBP2 dead E3 (hereafter referred to as RanBP2-dE3), where one copy of the gene (“f1”; Figure 3A-D) had a 45 base pair (bp) deletion just downstream from the targeted region (i.e., the guide RNA Protospacer Adjacent Motif “PAM” site) that eliminated 15 amino acids in the SUMO E3-ligase domain, and where the second copy (“f2”; Figure 3A-D) had a 356 bp deletion which eliminated the remaining part of exon 21 and a portion of the following intronic sequence (Figure 3A). We reasoned that the splicing of mRNAs generated from the f2 copy would produce a mature mRNA that contains premature stop codons and thus should be eliminated by nonsense-mediated decay (NMD). Thus, we inferred that the cell line only produces protein from the f1 copy of the gene, which should lack a portion of the E3 domain. The E3 domain not only binds to Ubc9, which is the only known SUMO E2-ligase in humans, but also to SUMO-RanGAP1 (Saitoh et al., 1997). In agreement with this, the nuclear rim localization of RanGAP1 was disrupted in RanBP2-dE3 cells (Figure 3E) despite the fact that the mutant protein was still at the nuclear rim (Figure S2). RanBP2-dE3 cells also had significantly decreased levels of RanBP2 protein, although this appeared to vary greatly between experiments (for example compare levels of RanBP2 in RanBP2-dE3 cells in Figure 3F and H), and had almost no detectable SUMO-conjugated RanGAP1 (SUMO-RanGAP1) (Figure 3F, 3H). Importantly, when RanBP2-dE3 cells were transfected with *IL6-Δi-HA* or *IL6-1i-HA*, these cells had significantly elevated IL6-HA protein expression over control cells (Figure 3F-I). Note that co-transfected Histone1-GFP (“H1B-GFP”) expression was similar in both cell lines (Figure 3F, 3H) indicating that there wasn’t a general alteration in mRNA translation. Confirming that the effect on IL6 expression was due to a decrease in translation, we fractionated lysates from unmodified and RanBP2-dE3 U2OS cells on a sucrose gradient (Figure 3J) and assessed the distribution of mRNA by RT-qPCR (Figure 3K-L). We found that *IL6-HA* mRNA was shifted towards the polysome fraction in mutant cells (Figure 3K). Meanwhile, the distribution of *α-tubulin* mRNA remained relatively unaffected (Figure 3L). These data indicate that either the SUMO E3-ligase activity, or high levels of RanBP2, is required for the translational repression of the *IL6* mRNA. We also examined the expression of TNF-α, another ANE1-associated cytokine (Table S1). As with IL6, the level of TNF-α-HA protein increased in RanBP2-dE3 cells (Figure 3M).

**Figure 3.**
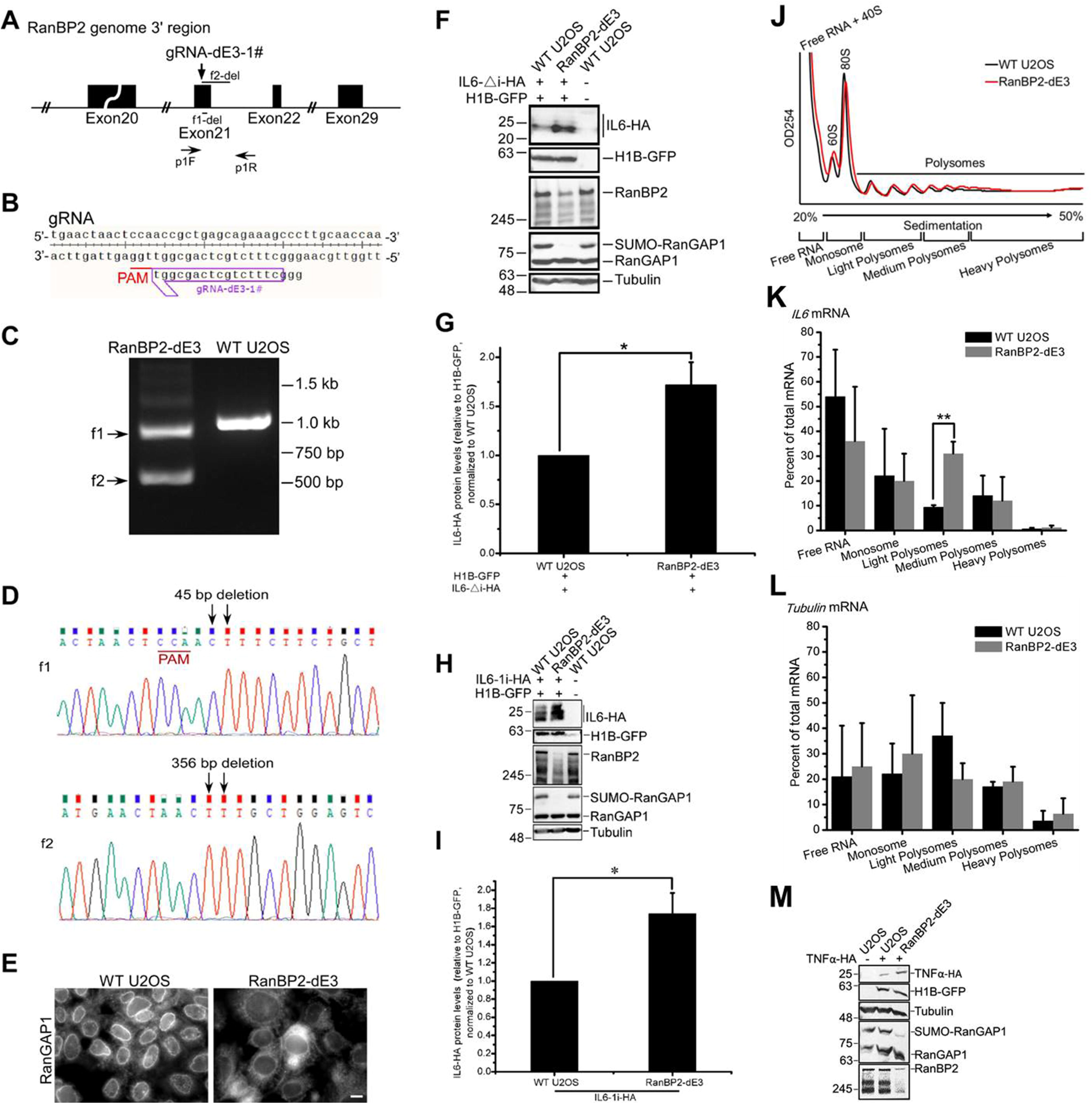
The SUMO E3-ligase domain of RanBP2 is required for IL6 translational suppression in U2OS cells. The SUMO E3-ligase domain of RanBP2 was targeted by CRISPR/Cas9 in U2OS cells. (A) A schematic diagram of the region of the RanBP2 gene targeted by CRISPR/Cas9 loaded with the guide RNA, “gRNA-dE3-1#”, whose sequence is shown in (B). Also indicated are the PCR amplification primers “p1F” and “p1R”, and the regions deleted “f1-del” and “f2-del” in each *RanBP2* allele present in the cell clone “RanBP2-dE3”. (C) PCR amplification, using p1F and p1R primers, of the genomic region targeted by gRNA-dE3-1#. Note that the reaction from the RanBP2-dE3 cell clone lysates produced two amplicons (“f1” and “f2”), which were both smaller than the amplicon produced from unmodified wildtype U2OS cells (“WT U2OS”). (D) Sequencing of the two PCR products (f1 and f2) in (C). Note that the length of the deletion in each *RanBP2* gene allele are indicated. (E) WT U2OS and RanBP2-dE3 cells were fixed, immunostained using anti-RanGAP1 antibody and imaged by epifluorescence microscopy. Scale bar = 10 µm. (F-G) WT U2OS, and RanBP2-dE3 were transfected with plasmids containing an intronless version of *IL6-HA* (*IL6-Δi-HA*) and *histone 1B-GFP* (*H1B-GFP*). Cell lysates were collected 24 h post-transfection and separated by SDS-PAGE. Proteins were detected with by immunoblot with antibodies against HA, GFP, RanBP2, RanGAP1 and α-tubulin. Note that H1B-GFP was used as a control for transfection and general mRNA translation while α-tubulin was used as a loading control. Also note that RanBP2-dE3 cells had lower expression of RanBP2 and lacked sumoylated-RanGAP1. IL6-HA and H1B-GFP protein levels were quantified using densitometry analysis and the ratio of IL6-HA/H1B-GFP was normalized to WT U2OS cells. (H-I) Same as (F-G), except that an intron-containing *IL6-HA* construct (*IL6-1i-HA*) was used. (J-L) WT U2OS and RanBP2-dE3 cells were transfected with an intron-containing *IL6-HA* construct (*IL6-1i-HA*). 24 h later cells were lysed and fractionated by centrifugation over a sucrose gradient. (J) OD254 trace of the sucrose gradients to determine the distribution of monosomes and polysomes. RT-qPCR of *IL6* (K) and *α-tubulin* (L) mRNA normalized against *luciferase* mRNA that was spiked into each fraction to control for RNA recovery. Note the significant increase in *IL6* mRNA in the polysome fraction in RanBP2-dE3 cells. (M) As in (F) except that TNF-α-HA was transfected. Each bar is the average of three independent experiments ± SEM. \**P* = 0.01–0.05, \*\**P* = 0.001–0.01, (Student’s *t*-test).

To validate our findings, we also generated a RanBP2 sumoylation-deficient mutant in human HAP1 cells. As these are haploid, we only had to modify the single copy of the *RanBP2* gene. Using a similar strategy as described above for RanBP2-dE3 generation, we isolated a mutant HAP1 clone called RanBP2-E3 insertion mutant (hereafter referred to as RanBP2-E3ins), which had a 51 bp (17 amino acids) insertion into the middle of the E3 domain (Figure S3A-B). Although RanBP2-E3 ins had no effect on RanBP2 protein level, it eliminated RanGAP1 sumoylation (Figure S3C). As we had observed in U2OS cells, expression of IL6-HA protein from the *IL6-1i-HA* reporter was higher in RanBP2-E3 ins cells than control HAP1 cells, when normalized to H1B-GFP expression (Figure S3C-D).

Taken together, these results demonstrate that the SUMO-E3 domain of RanBP2 is required to repress IL6-HA reporter protein production in cells.

### RanBP2-mediated Translation Inhibition of the *IL6-HA* Reporter Requires Both the *IL6* 5′UTR and 3′UTRs

Next we dissected the *IL6* mRNA to identify RanBP2-responsive elements. Replacing the *IL6* SSCR with the *major histocompatibility complex* (*MHC*) SSCR, which promotes translation in a RanBP2-dependent manner (Mahadevan et al., 2013), did not affect RanBP2-dependent silencing (Figure S1D-F). In contrast, replacing either the *IL6* 5′UTR or the 3′UTR sequences within the *IL6-1i* construct with the corresponding sequences from *ftz* (to form *5F-IL6-1i* - 5′UTR swapped, and *IL6-1i-3F* - 3′UTR swapped; Figure 4A), disrupted RanBP2-dependent silencing (Figure 4B-C). Interestingly, the 5’UTR swapped construct produced very little protein in both control and RanBP2-depleted cells and this was validated when we swapped the 5 UTR with that of the human *β-globin* gene (to form the *5βG-IL6-1i* construct - Figure 4A; protein levels shown in Figure 4D-E). Thus, there appears to be multiple elements that are required for RanBP2-dependent silencing.

**Figure 4.**
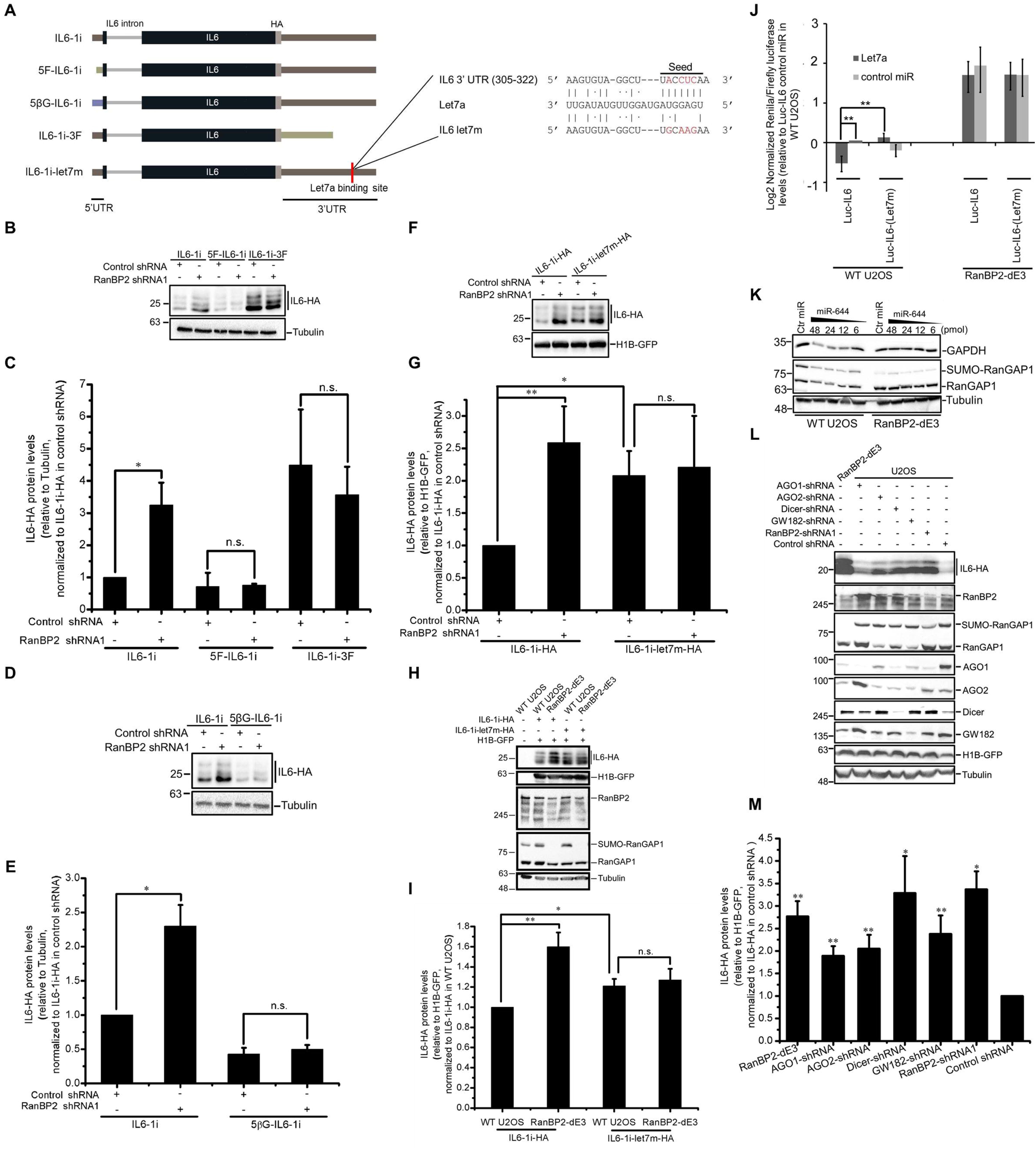
RanBP2 silences *IL6* mRNA through its *Let7*-binding site. (A) Schematic of the various intron-containing *IL6-HA* constructs (*IL6-1i*), where the 5′UTR was replaced with that of the *ftz* reporter (*5F-IL6-1i*) or the β-globin reporter (*5βG-IL6-1i*), and the 3′UTR was replaced with that of the ftz reporter (*IL6-1i-3F*) or had a mutation that eliminated the Let7 binding site (*IL6-1i-let7m*). (B-G) Cells treated with lentivirus that delivers shRNA1 against RanBP2, or scrambled shRNA (“control shRNA”) were transfected with the various *IL6-1i* constructs alone (B, D) or with H1B-GFP (F). 24 h post-transfection cell lysates were collected and separated by SDS-PAGE. The level of IL6-HA, α-tubulin and GFP were analyzed by immunoblot (B, D and F). The HA/tubulin (C,E) or the HA/GFP ratio (G) was quantified using densitometry analysis, normalized to *IL6-1i*-transfected control shRNA-treated cells, and plotted (C, E and G). (H-I) WT U2OS, and RanBP2-dE3 cells were transfected with *IL6-1i* or *IL6-1i-Let7m* and *H1B-GFP*. Cell lysates were collected 24 h post-transfection and separated by SDS-PAGE. Proteins were detected with by immunoblot with antibodies against HA, GFP, RanBP2, RanGAP1 and α-tubulin (H). IL6-HA and H1B-GFP protein levels were quantified using densitometry analysis and the ratio of IL6-HA/H1B-GFP was normalized to *IL6-1i-HA* transfected WT U2OS cells (I). (J) WT U2OS, and RanBP2-dE3 cells were co-transfected with *Let7a* miRNA or a scrambled control miRNA (“control miR”), and a dual luciferase plasmid that contains a *Renilla luciferase* reporter plasmid, carrying either the wild-type 3′UTR of *IL6* (*Luc-IL6*) or the *Let7a* binding site mutant (*Luc-IL6-(Let7m*), see Figure 7A for the sequence), and the Firefly *luciferase* as an internal control. 24 h after transfection, Renilla and Firefly luciferase luminescence were measured and the ratio was normalized to WT U2OS cells transfected with *Luc-IL6* and control miR. (K) WT U2OS, and RanBP2-dE3 cells were transfected with various amounts of miR-644 or 48 pmol of miR-144 (“Ctr miR”). 24 h post-transfection cell lysates were collected, separated by SDS-PAGE, and immunoprobed for GAPDH, RanGAP1 and α-tubulin. (L-M) WT U2OS cells treated with lentivirus that deliver shRNA against various proteins, or scrambled shRNA (“control shRNA”) were transfected with plasmid containing the *IL6-1i* construct. For comparison RanBP2-dE3 cells were also included in this analysis. Cell lysates were collected 24 h post-transfection and separated by SDS-PAGE. Proteins were detected with by immunoblot with antibodies against HA, RanBP2, RanGAP1, AGO1, AGO2, Dicer, GW182, GFP and α-tubulin (L). IL6-HA and H1B-GFP protein levels were quantified using densitometry analysis and the ratio of IL6-HA/H1B-GFP was normalized to *IL6-1i-HA* transfected WT U2OS cells (M). Each bar is the average of 3 independent experiments ± SEM. \**P* = 0.01–0.05, \*\**P* = 0.001–0.01, n.s. indicates no significant difference (Student’s *t*-test).

### The *Let7* Binding Site within the *IL6* 3′UTR Is Required for RanBP2-supression of IL6 Protein Production

Next, we decided to characterize the 3’UTR element RanBP2-responsive element. We found that *IL6-1i-3del1,* which only contains the first 110 nucleotides of the IL6 3’UTR (Figure S4A), was still regulated by RanBP2 (Figure S4B-C), while *IL6-1i-3del2*, which contains the last 317 nucleotides (Figure S4A), was no longer regulated by RanBP2 (Figure S4B-C). We noted that this deletion eliminated a *Let7a* miRNA binding site. Previously it was reported that *Let7a* miRNA directly inhibits expression of IL6 through this site (Iliopoulos et al., 2009; Schulte et al., 2011), and that RanBP2 is required for *Let7a*-mediated translation suppression (Sahoo et al., 2017). Importantly, several members of the *Let7* family of miRNAs are expressed in U2OS cells (Liu et al., 2014; Sohn et al., 2012). To determine whether the *Let7* binding site is required for RanBP2-mediated suppression of IL6, we generated a mutant of the *IL6-1i-HA* reporter bearing 4 point mutations in the *Let7* recognition site (*IL6-1i-Let7m-HA*) (Figure 4A). As expected, the *IL6-1i-Let7m-HA* construct produced more protein compared to the *IL6-1i-HA* construct in control cells (Figure 4F-G). In contrast, the level of protein from *IL6-1i-Let7m-HA* was similar to *IL6-1i-HA* in RanBP2-depleted cells (Figure 4F-G). Similar results were obtained in RanBP2-dE3 cells (Figure 4H-I). Note that the level of protein generated from co-transfected H1B-GFP was similar in all the cell lines (Figure 4F, H).

To further confirm these results, we monitored the expression of Renilla *luciferase* reporter plasmids carrying the wild-type 3′UTR of *IL6* (*Luc-IL6*) or a mutant version that lacked the *Let7* binding site (*Luc-IL6-Let7m*) (Schulte et al., 2011) in cells co-transfected with miRNA mimics. These reporter plasmids also contained the Firefly *luciferase* gene, to control for general changes in gene expression. In control U2OS cells we observed that co-transfection of *Let7a* inhibited protein production from the *Luc-IL6* reporter, but did not impact expression from *Luc-IL6-(Let7m)* (Figure 4J), as reported by others (Schulte et al., 2011). Importantly, RanBP2-dE3 cells had significantly enhanced expression from the *Luc-IL6* construct, in comparison to the unmodified U2OS cells (Figure 4J). Moreover, in the RanBP2-dE3 cells introduction of *Let7a* did not repress *Luc-IL6* (Figure 4J).

These results indicate that RanBP2 represses IL6 expression in part through a *Let7*-mediated translational suppression of the *IL6* mRNA. The fact that the RanBP2-dE3 cells have an overall higher expression of all the *Luc-IL6* constructs, regardless of the presence of a *Let7* binding site, suggests that there may be other RanBP2-sensitive elements in the 3′UTR. It is also curious that the *Luc-IL6* and *Luc-IL6-(Let7m)* reporters are not differentially expressed (Figure 4J, compare *Luc-IL6* and *Luc-IL6-(Let7m)* with scrambled control miRNAs). This may be due to the fact that for the endogenous *Let7* to supress translation through the *IL6* 3′UTR, the *IL6* 5′UTR is required.

### RanBP-dE3 Cells Have a General miRNA-silencing Defect

Our previous results showing that RanBP2-dependent sumoylation led to a decrease in *Let7*-dependent silencing led us to ask whether these cells were compromised generally for miRNA activity. To determine whether the defect in *Let7* mediated silencing extends to other miRNAs, we transfected U2OS or RanBP2-dE3 cells with miR-644, which is known to repress the expression of GAPDH (Sikand et al., 2012). In agreement with the idea that the miRNAi is generally compromised in RanBP2-dE3 cells, the presence of miR-644 did not lead to a decrease in GAPDH levels in these cells, as it did in the wildtype U2OS cells (Figure 4K). It should be noted that at higher levels, miR-644 does inhibit GAPDH in the mutant cells (data not shown), suggesting that the miRNA pathway is not totally inoperative in these cells but is less active.

### RISC is Required to Silence IL6

Our data and previous results indicate that IL6 is regulated by the RNA induced silencing complex (RISC). To confirm this, we depleted various proteins in the RNAi pathway and monitored IL6-HA production. Depletion of AGO1, AGO2, dicer, and GW182 all led to an increase in IL6-HA protein production without affecting co-transfected H1B-GFP (Figure 4L-M). Interestingly, depletion of certain factors in this pathway led to changes in the levels of other factors suggesting that these proteins likely regulate each other. For example, the depletion of every RISC-associated protein and RanBP2 led to a precipitous drop in AGO1 levels (Figure 4L). AGO1 levels were also low in RanBP2-dE3 cells in comparison to unmodified U2OS. In contrast, depletion of AGO1 led to the upregulation of AGO2 protein levels (Figure 4L). To rule out that changes in AGO1 were due to IL6-HA production, we repeated these experiments in untransfected cells. Again, RanBP2-depletion or elimination of RanBP2-dependent sumoylation led to a drop in AGO1 levels but did not significantly affect the levels of AGO2 (Figure S5A-B).

In summary, our data suggested that not only was IL6 expression inhibited by RISC, but that RanBP2 may regulate IL6 by affecting AGO1 levels.

### RanBP2 Regulates IL6 Translation by Stabilizing of AGO1

Previously it had been reported that RanBP2 was required for the degradation of AGO2 (Sahin et al., 2014), however in other reports depletion of RanBP2 had no effect on AGO2 levels, but instead promoted the association of miRISC with its target mRNAs (Sahoo et al., 2017). Our new data suggested that RanBP2-dependent sumoylation was required for the stabilization of AGO1, but not AGO2. In agreement with this, treating RanBP2-dE3 cells with the proteasome inhibitor, MG132, increased AGO1 levels to that of WT U2OS cells (Figure 5A). This treatment also slightly increased AGO2 levels (Figure 5A). Interestingly, MG132-treatment also increased the levels of the mutant RanBP2 in RanBP2-dE3 cells (Figures 5A), suggesting that the 15 amino acid deletion destabilizes this protein. In contrast, MG132-treatment had no major effects on the levels of wildtype RanBP2, α-tubulin or RanGAP1. Remarkably, RanBP2-dE3 and wildtype cells treated with MG132 had much lower levels of IL6-HA than untreated cells (Figure 5A-B). Furthermore, MG132-treated RanBP2-dE3 and wildtype cells had similar levels of IL6-HA protein (Figure 5A-B), suggesting that when Argonaute proteins are prevented from degradation, RanBP2-dependent sumoylation was no longer required for IL6 suppression. Notably, this treatment did not alter global translation patterns as the levels of expression from other co-transfected reporters (H1B-GFP and Flag-HA-tagged EYFP) were similar across all cell lines and all treatments (Figure 5A).

**Figure 5.**
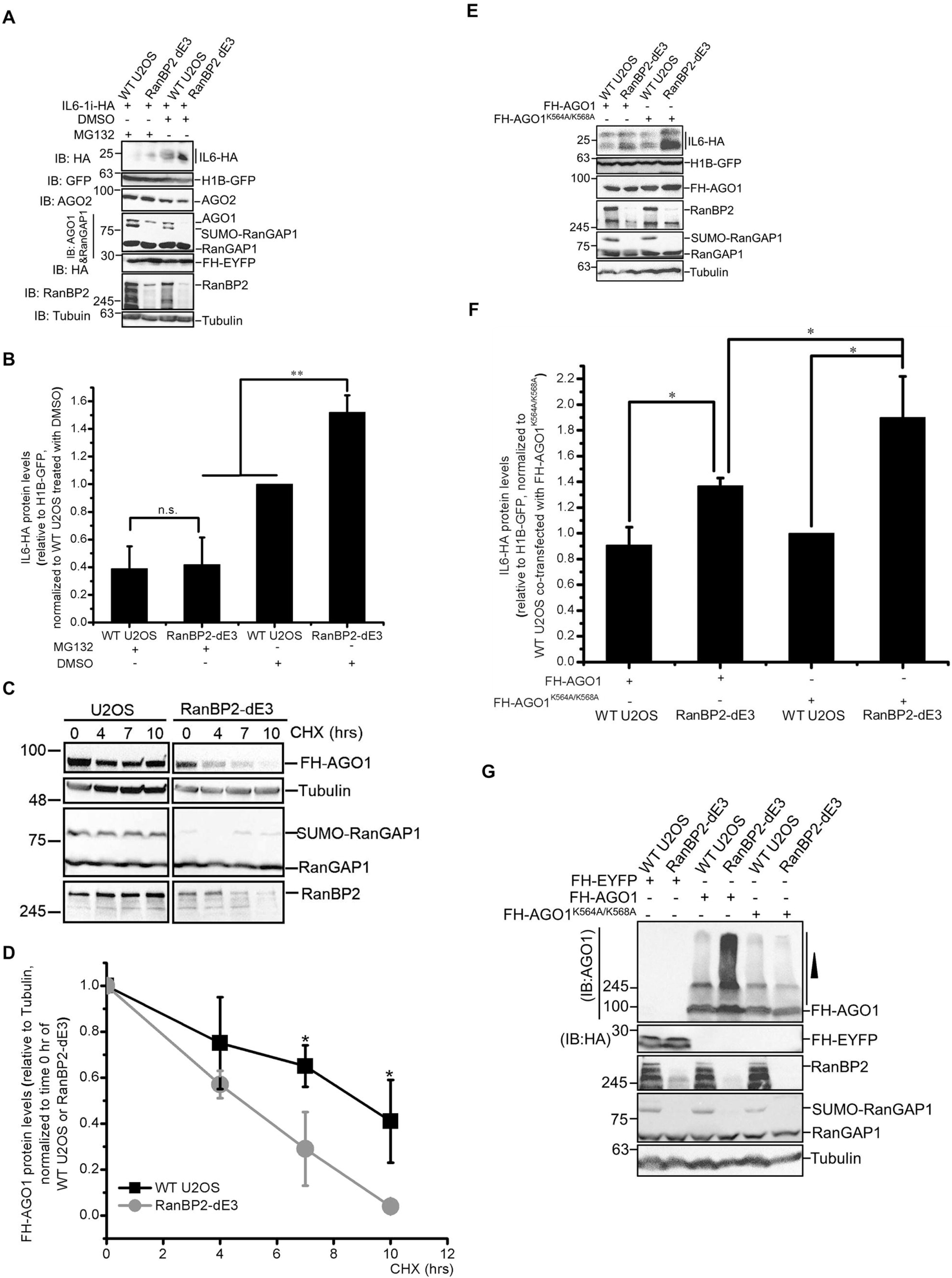
RanBP2 represses IL6 expression by promoting AGO1 stabilization. (A-B) WT U2OS, and RanBP2-dE3 cells were co-transfected with *IL6-1i-HA*, *H1B-GFP* and Flag-His-tagged yellow fluorescent protein (FH-EYFP). 18 h post-transfection, cells were treated with MG132 (10 µM) (MG132 “+”), or DMSO (MG132 “-”) for an additional 7 h. Cell lysates were collected, separated by SDS-PAGE, and immunoblotted with antibodies against HA, GFP, AGO2, AGO1, RanGAP1, RanBP2 and α-tubulin. Note that AGO1 and RanGAP1 are blotted together. Also note that DMSO-treated RanBP2-dE3 cells have no detectable AGO1. MG132-treatment led to an increase in AGO1, AGO2 and mutant RanBP2-dE3 levels. In contrast, the same treatment led to a decrease in IL6-HA levels. IL6-HA and H1B-GFP protein levels were quantified using densitometry analysis and the ratio of IL6-HA/H1B-GFP was normalized to DMSO-treated WT U2OS cells (B). (C-D) WT U2OS, and RanBP2-dE3 cells were transfected with Flag-HA-tagged AGO1 (FH-AGO1). 24 h post-transfection, the cells were treated with cycloheximide (CHX 100 µM) for various amounts of time to block further translation and thus allowing us to determine the rate of AGO1 turnover. Cell lysates were collected, separated by SDS-PAGE, and immunoblotted with antibodies against HA (FH-AGO1), α-tubulin, RanGAP1 and RanBP2 (C). For each time point, FH-AGO1 and α-tubulin protein levels were quantified using densitometry analysis and the ratio of FH-AGO1/ α-tubulin was normalized to the zero time point (D). (E-F) WT U2OS, and RanBP2-dE3 cells were co-transfected with *IL6-1i-HA*, *H1B-GFP* and either *FH-AGO1*, or *FH-AGO1^K564A/K568A^*. Cell lysates were collected 24 h post-transfection and separated by SDS-PAGE and immunoprobed for IL6-HA (anti HA), H1B-GFP (anti GFP), FH-AGO1 (anti HA), RanBP2, RanGAP1 and α-tubulin proteins (E). H1B-GFP was used as a control for transfection efficiency, and general mRNA translation. α-tubulin was used as a loading control. IL6-HA and H1B-GFP protein levels were quantified using densitometry analysis and the ratio of IL6-HA/H1B-GFP was normalized to FH-EYFP transfected WT U2OS cells (F). (G) WT U2OS, and RanBP2-dE3 cells were transfected with either *FH-AGO1*, *FH-AGO1^K564A/K568A^* or *FH-EYFP*. Cell lysates were collected 24 h post-transfection, separated by SDS-PAGE and immunoprobed for FH-AGO1, FH-EYFP, RanBP2, RanGAP1, and α-tubulin proteins. Each bar is the average of 3 independent experiments ± SEM. *\*P* = 0.01–0.05, *\*\*P* = 0.001–0.01, n.s. indicates no significant difference (Student’s *t*-test).

To confirm that AGO1 stability required RanBP2-dependent sumoylation, we treated cells with cycloheximide to prevent further expression of protein, and then assessed the rate at which the remaining AGO1 degraded. Since levels of AGO1 in RanBP2-dE3 cells were already very low, we monitored the levels of overexpressed Flag-HA-tagged AGO1 (FH-AGO1). Note that the presence of an N-terminal tag is known not to interfere with Argonaute-dependent silencing (Meister et al., 2004). Indeed, we observed that FH-AGO1 had a higher turnover rate in RanBP2-dE3 cells compared to unmodified U2OS cells (Figure 5C-D) and that inhibition of degradation with MG132 eliminated the difference in the levels of FH-AGO1 between these two cell lines (Figure S6).

Finally, we tested whether the lack of silencing in RanBP2-dE3 cells can be overcome by overexpressing AGO1. Indeed, overexpression of FH-AGO1 partially suppressed the expression of IL6-HA protein in these cells when compared to the expression of mutant AGO1 (FH-AGO1^K564A/K568A^) (Figure 5E-F), which does not bind to miRNAs (Boland et al., 2011). Expression of FH-AGO1 in control U2OS cells did not further reduce IL6-HA levels, suggesting that endogenous levels of AGO1 are sufficient to suppress IL6-HA expression (Figure 5E-F). Expression of FH-AGO1 had no effect on H1B-GFP expression (Figure 5E) indicating that general translation was not perturbed in these cells.

From these experiments we conclude that RanBP2-dependent sumoylation is required to stabilize AGO1, which in turn acts to suppress the translation of the *IL6-HA* reporter. This is in line with previous studies that have shown that the AGO1 protein strongly interacts with, and regulates the expression of *IL6* mRNA upon *Let7a* overexpression (Iliopoulos et al., 2009; Schulte et al., 2011), and that RanBP2 is required for the *Let7*-mediated suppression of *luciferase* reporter mRNAs (Figure 4J and (Sahoo et al., 2017)).

### RanBP2 Promotes the Sumoylation, and Inhibits the Ubiquitination, of AGO1

When analyzing the overexpression of various versions of FH-AGO1, we noted that RanBP2-dE3 cells accumulated huge amounts of high molecular weight AGO1 that were often confined to the stacking gel (Figure 5G). This suggested that a significant proportion of FH-AGO1 accumulated post-translational modifications in RanBP2-dE3 cells and raised the possibility that in the absence of RanBP2-dependent sumoylation, FH-AGO1 was poly-ubiquitinated and then targeted to the proteasome. This high mobility AGO1 was not seen with the FH-AGO1^K564A/K568A^ mutant (Figure 5G), which does not bind to miRNAs, suggesting that RanBP2 may only affect the post-translational modification of active AGO1.

To identify whether RanBP2 promotes the sumoylation of AGO1, we co-expressed FH-AGO1 with His-tagged SUMO2 (His6-SUMO2). To enhance sumoylation we co-transfected SV5-tagged Ubc9 (V5-Ubc9). Again, note that Ubc9 is the only known SUMO-conjugating E2 enzyme in humans. His6-SUMO2-conjugated FH-AGO1 was purified from denatured cell extracts on a nickel column, which binds to the His tag on the exogenous SUMO2, and the eluate was analyzed using an anti-HA antibody. As we suspected, FH-AGO1 was strongly sumoylated in wildtype U2OS and relatively weakly in RanBP2-dE3 cells (Figure 6A). No FH-AGO1 was seen in the nickel-bound fraction from cells that did not express His6-SUMO2, indicating that the signal was specific to His6-SUMO2-FH-AGO1. When total His-tagged SUMO2 conjugated proteins were assessed, we saw a modest decrease in RanBP2-dE3 relative to wildtype U2OS cells (Figure 6A).

**Figure 6.**
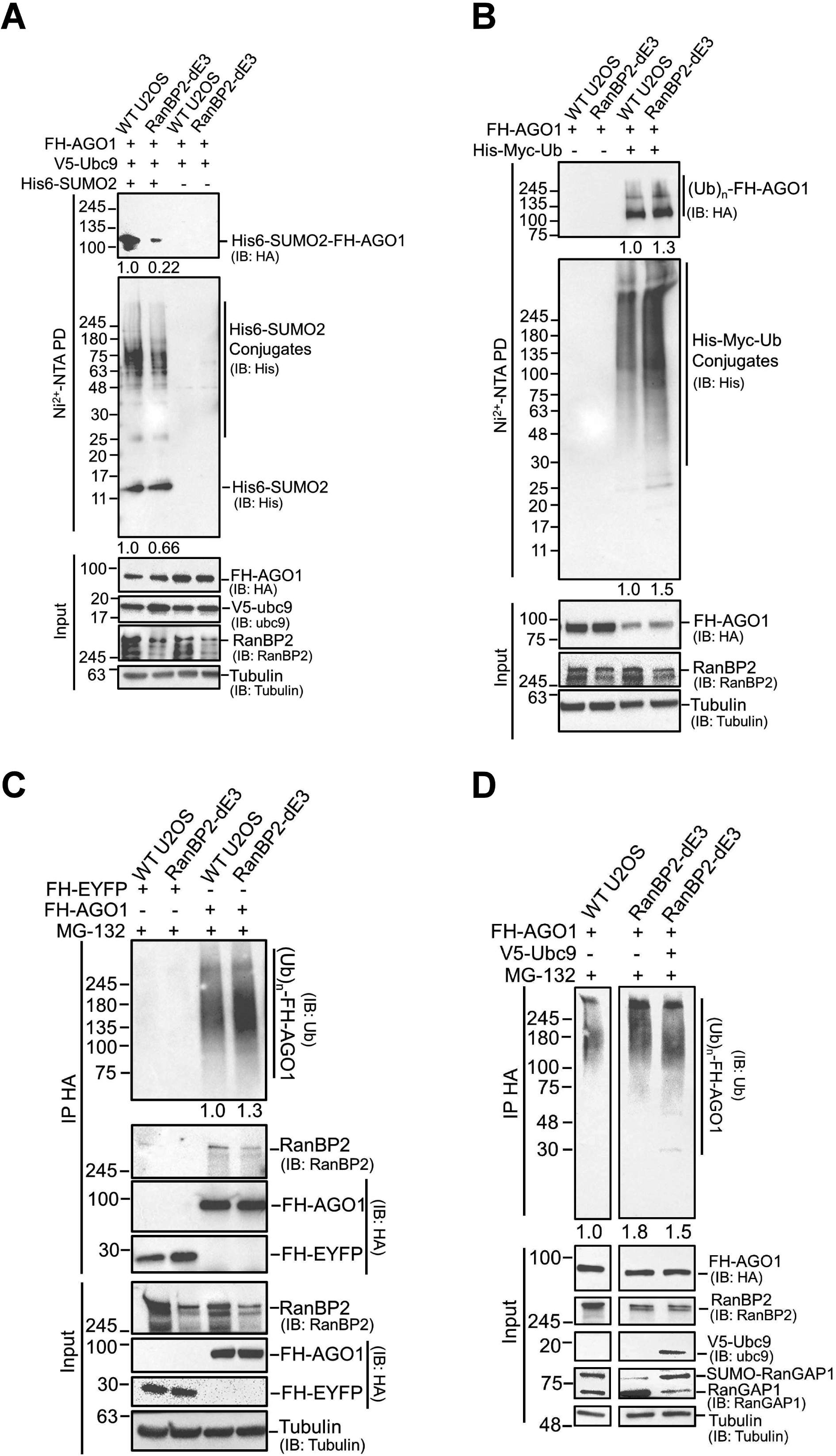
RanBP2 promotes the sumoylation and inhibits the ubiquitination of AGO1. (A) WT U2OS, and RanBP2-dE3 cells were co-transfected with *FH-AGO1*, *V5-Ubc9*, and His-tagged SUMO2 (*His6-SUMO2* “+”) or control vector (*His6-SUMO2* “-”). 24 h post-transfection cells were lysed in 6 M guanidinium chloride, and the His6-SUMO2 conjugates were isolated on Nickel beads (“Ni2+ NTA PD”) or the lysates were directly analyzed (“input”) and separated by SDS-PAGE. Conjugates were analyzed for the presence of FH-AGO1 by immunoblotting for HA (IB: HA), and for total His6-SUMO2 conjugates by immunoblotting for His (IB: His). Relative levels of each signal, as analyzed by densitometry, are indicated below each blot. Input lysates were immunoblotted for FH-AGO1, V5-Ubc9, RanBP2 and α-tubulin. (B) WT U2OS, and RanBP2-dE3 cells were co-transfected with FH-AGO1, and His-Myc-tagged ubiquitin (His-Myc-Ub “+”) or as V5-Ubc9 as a control (His-Myc-Ub “-”). 18 h post-transfection, cells were treated with MG132 (10 µM) for an additional 7 h to preserve ubiquitinated conjugates. Cells were lysed in 6 M guanidinium chloride, and the His-Myc-Ub conjugates were isolated on Nickel beads (“Ni2+ NTA PD”) or the lysates were directly analyzed (“input”) and separated by SDS-PAGE. Conjugates were analyzed for the presence of FH-AGO1 by immunoblotting for HA (IB: HA), and for total His-Myc-Ub conjugates by immunoblotting for His (IB: His). Relative levels of each signal, as analyzed by densitometry, are indicated below each blot. Input lysates were immunoblotted for FH-AGO1, RanBP2 and α-tubulin. (C) WT U2OS, and RanBP2-dE3 cells were transfected with FH-AGO1 or FH-EYFP. 18 h post-transfection, cells were treated with MG132 (10 µM) for an additional 7 h to preserve ubiquitinated conjugates. Cells were lysed in RIPA buffer, and the FH-AGO1/FH-EYFP and associated proteins were isolated by immunoprecipitation using anti-HA antibodies (“IP HA”) or the lysates were directly analyzed (“input”) and separated by SDS-PAGE. The immuoprecipitates were analyzed for ubiquitinated proteins by immunoblotting against ubiquitin (IB: Ub). They were also analyzed for the immunoprecipitated FH-AGO1/FH-EYFP by immunoblotting against HA, and for co-immunoprecipitated RanBP2. Input lysates were immunoblotted for RanBP2, FH-AGO1 and FH-EYFP (IB: HA) and α-tubulin. (D) WT U2OS, and RanBP2-dE3 cells were transfected with FH-AGO1 and either V5-Ubc9 to enhance sumoylation or control plasmid. 18 h post-transfection, cells were treated with MG132 (10 µM) for an additional 7 h to preserve ubiquitinated conjugates. Cells were lysed in RIPA buffer, and the FH-AGO1/FH-EYFP were isolated by immunoprecipitation using anti-HA antibodies (“IP HA”) or the lysates were directly analyzed (“input”) and separated by SDS-PAGE. The immuoprecipitates were analyzed for ubiquitinated proteins by immunoblotting against ubiquitin (IB: Ub). Input lysates were immunoblotted for FH-AGO1 (IB: HA), RanBP2, V5-Ubc9, RanGAP1 and α-tubulin. Note that Ubc9 overexpression, rescues RanGAP1-sumoylation in RanBP2-dE3 cells and decreases the amount of ubiquitinated FH-AGO1.(E) General model for the regulation of *IL6* by RanBP2.

Next, we wanted to assess the effect of the RanBP2 mutation on AGO1 ubiquitination. We co-expressed FH-AGO1 with His-Myc tagged ubiquitin (His-Myc-Ub) in wildtype and RanBP2-dE3 cells. To ensure that we would capture ubiquitinated intermediates, we inhibited protein degradation with MG132. We then purified ubiquitinated substrates from the cell lysates on a nickel column, which binds to the His-tag on the exogenously expressed ubiquitin. In contrast to what we had seen with sumoylation, the level of ubiquitinated FH-AGO1 was 30% higher in RanBP2-dE3 cells compared to wildtype U2OS (Figure 6B). Again, no FH-AGO1 was seen in the nickel-bound fraction from cells that did not express His-Myc-Ub, indicating that the signal was specific. Furthermore, when total ubiquitinated products were blotted for, using antibodies against the His tag, we saw a general increase in ubiquitinated substrates, indicating that the RanBP2-dE3 cells had higher levels of ubiquitinated proteins.

To confirm this last result, we immunoprecipitated exogenously expressed FH-AGO1 from MG132-treated cell lines and blotted for endogenous ubiquitin to visualize ubiquitinated conjugates. Again, the level of ubiquitinated AGO1 increased in RanBP2-dE3 cells by about 30% compared to wildtype U2OS (Figure 6C), even though equal amounts of FH-AGO1 were assessed (Figure 6C).

We next want to ensure that the reason that RanBP2-dE3 cells promoted an increase in ubiquitination was due to the loss of its ability to sumoylate downstream targets, and not some other activity that happened to be disabled by the RanBP2-dE3 mutations, such as binding to RanGAP1. To increase the overall levels of sumoylation in these cells, we overexpressed the V5-Ubc9. Indeed, V5-Ubc9 overexpression resulted in an increase in sumoylated RanGAP1 (Figure 6D), supporting the notion that this protocol did indeed result in an increase in overall sumoylation activity in RanBP2-dE3 cells (Figure 6D). When FH-AGO1 was immunoprecipitated and the level of its ubiquitin-conjugates was assessed by immunostaining, we observed that V5-Ubc9 overexpression reduced overall ubiquitination in RanBP2-dE3 cells (Figure 6D). Thus, the overexpression of Ubc9, and the accompanying general increase in sumoylation activity, reduced FH-AGO1 ubiquitination even in RanBP2-dE3 cells.

From these experiments we conclude that RanBP2 promotes the sumoylation of AGO1 and inhibits its ubiquitination. In addition, our experiments also suggest that sumoylation inhibits ubiquitination of AGO1. Although our experiments strongly indicate that AGO1 is a direct sumoylation substrate for RanBP2, we cannot rule out the possibility that some intermediate targets exist between RanBP2 and AGO1, though we believe that this is unlikely.

### RanBP2 Associates with AGO1

Previously it had been reported that RanBP2 directly interacts with AGO2 (Sahoo et al., 2017). Indeed, we found both RanBP2 and the RanBP2-dE3 mutant in immunoprecipitates of FH-AGO1 (Figure 6C), but not in control immunoprecipitates (FH-EYFP). This suggests that RanBP2 interacts with AGO1 in a manner analogous to AGO2. Interestingly, it was previously reported that this interaction required one of two putative SUMO-interacting motifs (SIMs) that are present in the E3 domain of RanBP2. Although the dE3 mutation deletes part of the first putative SIM, the second putative SIM is still intact. While this interaction would suggest that RanBP2 should bind to sumoylated Argonautes, Sahoo and colleages clearly detected interactions between RanBP2 and unmodified AGO2. Note that the interaction between RanBP2 and AGO1 further supports the notion that RanBP2 directly sumoylates Argonaute proteins.

### Expression of IL6 mRNA Promotes the Sumoylation of AGO1 and the Nuclear Export of RISC Components

Interestingly we observed that AGO1 and GW182 were both primarily localized to the nucleoplasm of U2OS cells (Figure 7A). Indeed various reports have indicated that endogenous RISC complexes are found in the nuclei of several tissue culture cell lines and in cells derived from various animal tissues (Ahlenstiel et al., 2012; Matsui et al., 2015; Rentschler et al., 2018; Robb et al., 2005; Sharma et al., 2016; Shuaib et al., 2019). The fact that at steady state most of the *IL6* mRNA is in the cytosol (Figure 2G), yet not being translated, must mean that AGO1 silences the cytoplasmic pool of mRNA. These observations raised the possibility that RISC is recruited to the *IL6* mRNA in the nucleus and that upon the completion of nuclear export, AGO1 gets sumoylated by RanBP2 as part of an mRNP maturation process. The sumoylation of AGO1 would then be required for its stable association to the *IL6* mRNA and thus promote *IL6* silencing.

**Figure 7.**
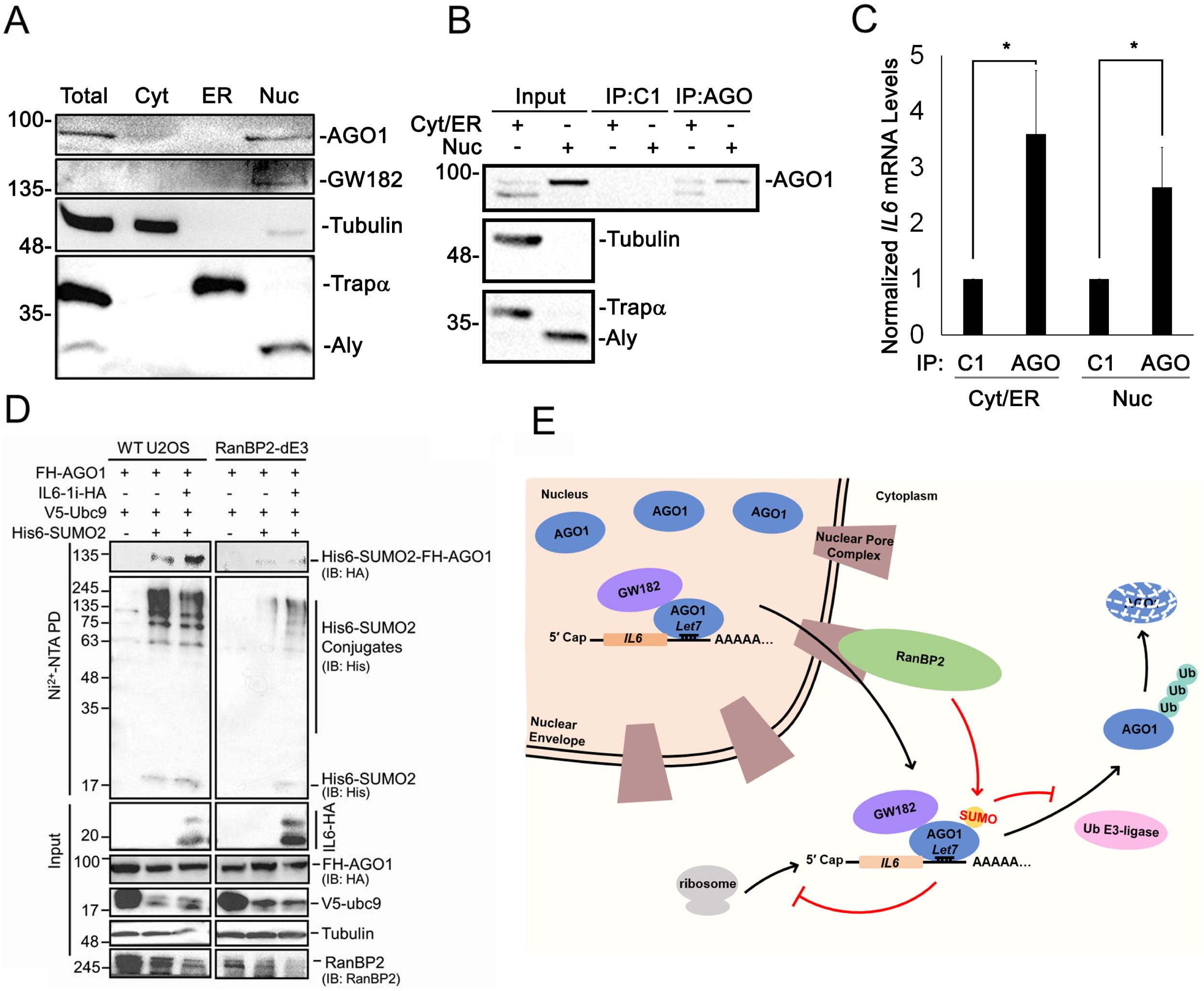
IL6 expression promotes the nuclear export of RISC and the RanBP2-dependent sumoylation of AGO1. (A) U2OS cells were fractionated into cytoplasmic, ER and nuclear fractions. Samples were separated by SDS-PAGE and immunoblotted for AGO1, GW182, tubulin (cytoplasmic marker), Trapα (ER marker) and Aly (nuclear marker). (B-C) Cells were transfected with *IL6-Δi-HA* then fractionated into cytoplasmic/ER and nuclear fractions. Fractions were either collected (“Input”) or immunoprecipitated with a control (C1) or anti-AGO synthetic antibody fragment. Samples were separated by SDS-PAGE and immunoblotted for AGO1, tubulin (cytoplasmic marker), Trapα (ER marker) and Aly (nuclear marker) (B). The amount of *IL6* mRNA in the immunoprecipitates was quantified by qRT-PCR, each bar being the average ± SEM of 4 experiments (C) \**P* = 0.01–0.05. (D) WT U2OS, and RanBP2-dE3 cells were co-transfected with *FH-AGO1*, *V5-Ubc9*, with or without His-tagged SUMO2 (*His6-SUMO2* “+/-”) and *IL6-i-HA* (“+/-“). 24 h post-transfection cells were lysed in 6 M guanidinium chloride, and the His6-SUMO2 conjugates were isolated on Nickel beads (“Ni2+ NTA PD”) or the lysates were directly analyzed (“input”) and separated by SDS-PAGE. Conjugates were analyzed for the presence of FH-AGO1 by immunoblotting for HA (IB: HA), and for total His6-SUMO2 conjugates by immunoblotting for His (IB: His). (E) General model for how RanBP2 regulates the silencing of the *IL6* mRNA through the sumoylation of AGO1.

To test this model we wanted to determine whether nuclear AGO1 could associate with *IL6* mRNA. We thus isolated cytosolic and nuclear fractions from cells expressing *IL6-Δi-HA* (Figure 7B), and immunoprecipitated endogenous AGO1 using a synthetic antibody (Na et al., 2016) and tested for the presence of *IL6* mRNA by RT-qPCR. As predicted by our model, *IL6* mRNA was enriched in the immunoprecipitated nuclear AGO1 fraction (Figure 7C).

Finally, our model predicts that the over-expression of *IL6* mRNA should drive more RISC through the nuclear pore and hence increase the sumoylation of AGO1. In agreement with this we observed that U2OS cells expressing IL6 have higher levels of sumoylated FH-AGO1 (Figure 7D). This was dependent on RanBP2, as the increase in sumoylation was not seen in RanBP2-dE3 cells.

In summary our data supports our model that in U2OS cells, RISC is loaded onto the *IL6* mRNA in the nucleus, and that upon the completion of nuclear export, AGO1 is sumoylated by RanBP2. This post-translational modification likely stabilizes AGO1 onto the mRNA thus enforcing silencing (Figure 7E).

## DISCUSSION

In this study, we demonstrate that the nuclear pore filament protein RanBP2, sumoylates the *IL6* mRNP component AGO1, which stabilizes it and ultimately promotes the silencing of the *IL6* mRNA. Our data is consistent with the idea that this sumoylation event happens in conjunction with nuclear export of the *IL6* mRNP complex. First, we show that AGO1 associates with *IL6* mRNA in the nucleus. Next, we show that the over-expression of *IL6* mRNA drives the sumoylation of AGO1, indicating that once AGO1 joins the *IL6* mRNP it becomes a substrate for RanBP2. Finally, it is clear from our data that *IL6* mRNA is translationally repressed in the cytoplasm in a manner that requires RanBP2-dependent sumoylation (Figure 3J-K), and not targeted for decay (Figure 2E-F), an activity that has been document for nuclear argonautes (Gagnon et al., 2014). According to our model, RISC components that accompany mRNAs from the nucleus to the cytoplasm, but fail to be sumoylated, are targeted for ubiquitination and subsequent degradation. This model is consistent with our finding that the FH-AGO1^K564A/K568A^ mutant, which lacks miRNA binding and does not attach to mRNAs, does not appear to be subjected to this type of regulation (Figure 5G). We suspect that sumoylation may stabilize AGO1 within the mRNP and a lack of this modification may eject AGO1, leading to its subsequent ubiquitination and degradation. Alternatively, ubiquitination may itself trigger AGO1 dissociation from the mRNA. It is possible that AGO2 may also regulate the *IL6* mRNA in a similar manner, however it does not require RanBP2-dependent sumoylation for its stability. It is possible that AGO2 simply requires RanBP2-dependent sumoylation for its association to the mRNP and that the other subsequent steps differ with what we’ve documented for AGO1.

We cannot rule out the possibility that AGO1 initiates interactions with *IL6* mRNA in the cytoplasm, and that this mRNP then travels to the nuclear pore to become sumoylated by RanBP2, however we think that this is unlikely. miRNA-silenced mRNPs have not been documented to travel from sites within the cytoplasm to the nuclear pore. In light of this, we believe that RanBP2 must sumoylate AGO1 as it crosses the pore as part of an mRNP.

Our study is also one of the first to address the role of RanBP2 in the regulation of ANE1-associated cytokines. Until now it remained unclear how the ANE1-associated mutations in RanBP2 might contribute to pathology. Our results indicate that one possible contributing factor is that RanBP2 regulates the expression of two ANE1-associated cytokines, IL6 and TNF-α. In agreement with our findings, it is well known that *Let7* is a major regulator of the *IL6* mRNA (Iliopoulos et al., 2009; Schulte et al., 2011) and that this miRNA regulates inflammation signalling (Brennan et al., 2017; Lin et al., 2017). Moreover, there is evidence that *Let7* levels change in response to infection (Ma et al., 2012; Makkoch et al., 2016; Mazumder et al., 2013; Schulte et al., 2011). From all this, an overall model emerges where upon infection, *Let7* and RanBP2 modulate the inflammation response by downregulating IL6, and potentially other cytokines. A defect in this regulation may cause a hyperinflammatory response that leads to pathology.

Given all this, it remains unclear how ANE1 mutations may affect RanBP2 regulation of the RISC complex. Since the 5’UTR of the *IL6* mRNA is also required for RanBP2-dependent silencing, it is possible that RanBP2 regulates several different aspects of mRNP maturation besides RISC-mediated silencing. Our data also indicates that other *cis*-elements may be present in the 3′UTR. We inferred this by the fact that out *luciferase* reporter that contains the *IL6* 3′UTR was upregulated in the RanBP2-dE3 cells compared to unmodified cells and this was independent of the *Let7* binding site. Thus, RanBP2 likely regulates the translation of *IL6* mRNA through an additional element in the 3′UTR. This may also explain why AGO1 overexpression does not fully inhibit IL6 production in RanBP2-dE3 cells (Figure 6G-H). Other *cis-* and *trans*-factors that regulate the *IL6* mRNA are known (Higa et al., 2018; Iwasaki et al., 2011; Masuda et al., 2013; Mino et al., 2015; Muller et al., 2015), however it is unclear if these are responsive to RanBP2. These elements and factors all regulate *IL6* mRNA stability, so their connection to RanBP2-based regulation seems unlikely. Future studies will be needed to identify these supplementary elements (in the 5′ and 3′UTR) and their modes of action. Since ANE1 mutants are only 40% penetrant, other host factors may enhance or disrupt interactions between the mutated region of RanBP2 and modulators of *IL6* mRNA. ANE1 mutations may also affect some other activity of RanBP2. If this is true, RanBP2 may coordinate several processes that impact the immune response.

Interestingly, many small RNA pathways (i.e. germ granule small RNAs in *Caenorhabditis elegans* and piRNAs in *Drosophila*, zebrafish and mouse) involve the processing and/or loading of small RNAs onto their target complexes in large phase-separated structures, called nuages or germ granules, that are physically associated with nuclear pores (Voronina et al., 2011). Indeed, a recent report demonstrated that RanBP2 was required for piRNA silencing of transposable elements in *Drosophila* (Parikh et al., 2018). These studies lend support to our model that RanBP2 may help to assemble or simply stabilize a repressive complex (i.e. the RISC complex in this case) onto the *IL6* mRNA, after it emerges from the nuclear pore.

Our model indicates that, in at least certain cell lines, RISC complexes associate with their target mRNAs in the nucleus. This may depend on the subcellular localization of RISC components in various cell lines. Interestingly we have observed that N-terminally tagged AGO1 is about 70% cytoplasmic in U2OS cells by both immunofluorescence and cell fractionation. In contrast, over-expressed untagged AGO1 is almost completely nuclear. This suggests that the N-terminal tag partially disrupts nuclear localization. Interestingly, tagged versions of argonautes are extensively use throughout the field as most antibodies against the untagged proteins do not work well for immunoprecipitations or immunofluorescence. Thus it is likely that the degree to which RISC complexes are distributed to the nucleus may be underappreciated by many investigators.

Argonautes are known to be extensively post-translationally modified, especially by ubiquitination (Baumberger et al., 2007; Bronevetsky et al., 2013; Chinen and Lei, 2017; Derrien and Genschik, 2014; Josa-Prado et al., 2015; Kobayashi et al., 2018; Nayak et al., 2018; Sahin et al., 2014). Our data suggests that RanBP2 directly sumoylates AGO1, as it has been previously reported for AGO2 (Josa-Prado et al., 2015; Sahin et al., 2014; Sahoo et al., 2017), and that this sumoylation antagonizes AGO1 ubiquitination. How this would work is unclear at the moment. Sumoylation of a particular lysine residue may prevent the same residue from being ubiquitinated and thus stabilize the protein, as reported in other cases (Desterro et al., 1998). Alternatively, sumoylation of AGO1 may mask the binding site of an E3 ubiquitin-ligase or help recruit a ubiquitin protease. We however cannot exclude the possibility that additional sumoylated proteins are required to stabilize AGO1. Previously, it has been reported that sumoylation promotes AGO2 degradation (Josa-Prado et al., 2015; Sahin et al., 2014), although other reports have suggested that sumoylation is required to load miRNAs into AGO2 (Sahoo et al., 2017). Interestingly, the ubiquitination site of *Drosophila* AGO1 (the homolog of human AGO2), K514, which is recognized by the RING-type E3 ubiquitin ligase Iruka (Kobayashi et al., 2018), is not conserved within human AGO1, suggesting that AGO1 and AGO2 may have different modes of regulation. These possibilities can be sorted out by mapping the relevant sumoylated and ubiquitinated residues on AGO1 and determining whether mutating these residues disrupts RanBP2 dependent regulation. Uncovering the E3 ubiquitin-ligase would help to clarify these issues.

## MATERIALS AND METHODS

### Plasmid constructs

For all expression experiments, pCDNA3.1 plasmid containing the human *insulin* cDNA (Palazzo et al., 2007) and human *β-globin* (Akef et al., 2013); pEGFP plasmid containing the *H1B-GFP* fusion gene (Contreras et al., 2003) were described previously. The *interleukin 6* (*IL6*) gene cloned inside pSPORT6 plasmid was purchased from OpenBiosystems, and various versions of the *IL6* gene including *MHC-IL6-Δi, IL6-1f, IL6-1i, 5F-IL6-1i, 5βG-IL6-1i, IL6-1i-3F, IL6-1i-3del1, IL6-1i-3del2,* and *IL6-1i-Let7m*, were cloned inside the pcDNA 3.1 plasmid and associated mutations were made by restriction-enzyme cloning or site-directed mutagenesis (according to manufacturer’s protocol). pIRESneo-*FLAG/HA AGO1* (Addgene plasmid # 10820, FH-AGO1) and pIRESneo-*FLAG/HA EYFP* (Addgene plasmid # 10825, FH-EYFP) were gifts from T. Tuschl (Meister et al., 2004). The AGO1 site mutant construct FH-AGO1^K564A/568A^, was constructed from FH-AGO1 by restriction-enzyme-free cloning (van den Ent and Löwe, 2006) using the following primer sequences, K564A/568A-forward primer: ATGTCGCACTTGGTGGCATTAACAACATCCTAG and K564A/568A-reverse primer: TGATCGCGAGGCAGAGGTTGGACAGAGTCTG. The human *His6-SUMO2* (in pcDNA3), *V5-Ubc9* plasmid and *His-Myc-ubiquitin* plasmid were gifts from L. Frappier (De La Cruz-Herrera et al., 2018; Li et al., 2010; Vertegaal et al., 2006). shRNA plasmids (details below) were purchased from Sigma; the *CRISPR/CAS9* plasmid, pSpCas9(BB)-2A-Puro (PX459) V2.0 was a gift from F. Zhang (Addgene plasmid#62988) (Ran et al., 2013), and plasmids of Dual Luciferase Reporter Assay (psiCHECK2-Luc-IL6 and psiCHECK2-Luc-IL6-(Let7m), details below) were gifts from J. Vogel (Schulte et al., 2011). TNF-alpha (ORF and UTR) was first amplified from U2OS genomic cDNA and then cloned into pcDNA3. The HA tag was then inserted at the 3’ end of the ORF by restriction enzyme free cloning.

### Cell culture, cell transfection and lentiviral mediated shRNA knockdown

Cell culture and transfection were carried out as described previously (Gueroussov et al., 2010; Mahadevan et al., 2013). Briefly, both human osteosarcoma (U2OS) and embryonic kidney 293T (HEK293T) were maintained in Dulbeco’s Modified Eagle Medium (DMEM) supplemented with 10% fetal bovine serum, and 1% penicillin-streptomycin (WISENT). HAP1 cells (a gift from Alexio Muise) can be obtained from Horizon Genomics (Vienna, Austria). HAP1 cells were grown in Iscove’s modified Dulbecco’s medium (IMDM) supplemented with 10% fetal calf serum and 1% penicillin-streptomycin (WISENT). All cells were cultured at 37 □ in a 5% CO_2_-humidified incubator. For chemical treatments, MG132 (Sigma) and cycloheximide (CHX) (Sigma) were dissolved in DMSO and used at a final concentration of 10 µM and 100 µM, respectively.

All cells were plated 24 h before transfection and transfected at a confluency of 70-80% using GenJet-OS in vitro transfection reagent for U2OS cells (SignaGen Laboratories, Gaithersburg, MD, USA) or JetPRIME for 293T cells (PolyPlus) or Turbofectin for HAP1 cells (OriGene), following the manufacturer’s protocol.

Lentiviral-mediated shRNA knockdown was carried out as described previously (Cui et al., 2012) with plasmids encoding shRNA against RanBP2 (shRNA1: TRCN0000003452, shRNA3: TRCN0000003454, Sigma), AGO1 (shRNA: TRCN0000007859, Sigma), AGO2 (shRNA: TRCN0000011203, Sigma), Dicer (shRNA: TRCN0000051258, Sigma), GW182 (shRNA: TRCN0000376423, Sigma), or control vector (pLK0.1). Briefly, plasmids encoding shRNA were transfected into the HEK293T cells together with the accessory plasmids, VSVG and Δ8.9, to generate lentivirus carrying specific shRNA plasmids. Lentivirus was harvested from the medium 48 h post-transfection by filtering through a 0.44 μm filter. For infection, lentivirus was applied to U2OS cells with 8 µg/ml hexamethrine bromide. Puromycin was applied to the cell 24 h post-infection at 2 µg/ml to select for infected cells, and puromycin containing medium was changed every other day. Cell lysates were collected 5 days post-infection to assess the level of knockdown, and the cells were used for various experiments as described.

### Immunoblotting and immunoprecipitation

For immunoblotting, various culture cell lines were lysed with lysis buffer containing 50 mM Tris-HCl, 150 mM NaCl, 1% Triton X-100, 1 mM EDTA, and complete protease inhibitor cocktail (Roche), pH 7.4, on ice for 30 min. For immunoprecipitation, whole-cell extracts were collected 24-48 h after transfection and lysed in lysis buffer on ice for 30 min. After centrifugation for 30 min at 13,000 *g*, 4 °C, supernatants were collected and incubated with Protein-G Sepharose beads coupled to specific antibodies (2 μg per pulldown) for 2-3 h with rotation at 4 °C. The beads were washed 3 times with lysis buffer and bound proteins were eluted by boiling for 10 min in sample buffer containing 50 mM Tris-HCl (pH 6.8), 2% SDS, 10% glycerol, 0.1% bromophenol blue and 1% β-mercaptoethanol. For immunoblot analysis, immunoprecipitates or whole-cell lysates were separated by SDS-PAGE, transferred to nitrocellulose membrane and probed with primary antibodies against HA (HA-7 mouse monoclonal, 1:2000 dilution, Sigma; or rabbit polyclonal, 1:1000 dilution, Sigma), α-tubulin (mouse monoclonalDM1A, 1:1,000 dilution, Sigma), GFP (rabbit polyclonal, 1:1000 dilution, Invitrogen), RanGAP1 (mouse monoclonal, 1:1000 dilution, Santa Cruz), RanBP2 (mouse monoclonal mAb414 1:5000, Cederlane, or rabbit polyclonal, 1:1000 dilution, Abcam), Ubc9 (rabbit polyclonal, 1:1000, Cell Signaling), IL6 (mouse monoclonal, 1:2000 dilution, Abcam), AGO2 (11A9 rat monoclonal, 1:1000 dilution, Millipore), His (mouse monoclonal, 1:1000 dilution, Abcam), or AGO1 (rabbit monoclonal, 1:1000 dilution, Cell Signalling), ubiquitin (rabbit polyclonal, 1:500 dilution, Dako), or GAPDH (rabbit polyclonal, 1:1000 dilution, ABGENT), GW182 (rabbit polyclonal, 1:1000 dilution, Abcam), Dicer (rabbit polyclonal, 1:1000 dilution, Cell Signaling). Subsequently, the relevant horse radish peroxidase (HRP) conjugated anti-rabbit (1:2000, Cell Signaling) or anti-mouse (1:4000, Cell Signaling) secondary antibody was used. Chemiluminescence luminol reagent (Pierce) and the Versadoc system (Bio-Rad) were used to visualize the blots. ImageJ (NIH) was used for densitometry analysis.

### Fluorescent In Situ Hybridization (FISH) and immunofluorescence microscopy

Fluorescence in situ hybridization (FISH) staining was done using DNA specific probes against *IL6* (GTAACATGTGTGAAAGCAGCAAAGAGGCACTGGCAGAAAACAACCTGAAC, 5′ labelled with Alexa546, IDT) at a dilution of 1:500 in 60% formamide hybridization buffer as previously described (Gueroussov et al., 2010). Samples were mounted on Fluoromount with 4’,6-diamidino-2-phenylindole (DAPI) (Southern Biotechnologies, Birmingham, AL, USA). Immunofluorescence staining was performed as previously described (Cui et al., 2012; Gueroussov et al., 2010) using antibody against RanGAP1 (mouse monoclonal, 1:250 dilution, Santa Cruz) and a secondary antibody (Alexa647-conjugated donkey anti-mouse polyclonal; 1:1000; Life Technologies, Carlsbad, CA, USA). Microscopy, imaging and nuclear mRNA export quantifications were performed as previously described (Gueroussov et al., 2010; Mahadevan et al., 2013; Palazzo et al., 2007). An epifluorescence microscope on a TI-E inverted Nikon microscope using a 60X phase 2, oil objective and a Coolsnap HQ2 14 bit CCD camera (photometric, Tucson, AZ, USA) controlled using NIS elements Basic Research Microscope Imaging Software (2009) was used to capture all the images. Image exposures varied from 30 ms to 2 s. Data pertaining to total integrated intensity, cytoplasmic/total, and nuclear/total fluorescence intensity was calculated as previously described (Gueroussov et al., 2010) from raw, unprocessed images. Images shown in figures were adjusted for brightness and contrast using the Photoshop (Adobe).

### RNA isolation and Northern blotting

After 18–24 h of transfection, total RNA from cultured cells was extracted with Trizol Reagent (Invitrogen) according to the manufacturer’s protocol. RNA was separated on a denaturing agarose gel, transferred, and probed for *IL6* and *Tubulin* as previously described (Mahadevan et al., 2013).

### Generation of the SUMO E3 domain mutants of RanBP2 by CRISPR/Cas9

Genome editing of RanBP2 in U2OS or HAP1 cells was performed as previously described (Moyer and Holland, 2015). In brief, gRNA (5′-GGGCTTTCTGCTCAGCGGT-3′) that targets exon 21 of RanBP2 (Figures 3A, 4A) was inserted into PX459 V2.0 plasmid and transfected into U2OS and HAP1 cells with GenJet-OS (SignaGen) and Turbofectin (Origene) respectively according to the manufacturer’s instructions. After 48 h of transfection, cells were subjected to transient selection with 2 µg/ml puromycin for 1-2 days to enrich for transfected cells. After puromycin selection, single colonies were grown for 2-4 weeks and then part of the colony was subjected to genomic DNA extraction with lysis buffer (10 mM Tris-HCl, pH 7.5, 10 mM of ethylene diamine tetra-acetic acid, 0.5% of sodium dodecyl sulfate, 10 mM of NaCl, 1 mg/mL of proteinase K) as previously described (Moyer and Holland, 2015). PCR screenings were done using p1F and p1R (p1F sequence: 5′-GAACACTAAATCAGGATGCTAATTCTAG-3′ and p1R sequence: 5′-TCTCTTTCTGCTAGAGCTTTAGCTC-3′) primers to assess for presence of insertions and deletions (Indels) that had been made in the region of interest. Any positive hits from the PCR screening were further verified by Sanger sequencing. In addition, part of the colony was harvested for western blotting and immunofluorescence analysis of localization and post-translational modification status of RanGAP1 as described in the previous sections.

### Dual luciferase reporter assay

For the *IL6* 3′UTR luciferase assay, U2OS and RanBP2-dE3 cells (4×104 cells/well) were seeded in 24-well plates 16 h prior to transfection and transfected with psiCHECK2-Luc-IL6 (100 ng) or psiCHECK2-Luc-IL6-(Let7m) (100 ng) together with 50 nM has-Let7a-5p miRNA (MIMAT0000062, Dharmacon) or miRIDIAN microRNA Mimic Negative Control #1 (CN-001000-01-05, Dharmacon) using JetPRIME reagent (PolyPlus) according to the manufacturer’s instructions. Cells were collected 24 h after transfection and luciferase activities were measured with Dual-Luciferase Reporter Assay System (Promega) according to manufacturer’s instructions.

### Sumoylation and ubiquitylation assay

In vivo AGO1 sumoylation was analysed in U2OS and RanBP2-dE3 cells as previously described (De La Cruz-Herrera et al., 2018; Shen et al., 2017; Tatham et al., 2009). Briefly, U2OS and RanBP2-dE3 cells in 10 cm dishes were transfected with plasmids (2.5 μg each) expressing His6-SUMO2 and V5-Ubc9, together with FLAG/HA-tagged AGO1 (FH-AGO1) proteins (in pIRESneo) or empty plasmid negative control using JetPRIME reagent (PolyPlus) according to the manufacturer’s instructions. 24 h after transfection, the cells were harvested, and 10% were lysed in 2X SDS loading buffer (60 mM Tris-HCl pH 6.8, 1% SDS, 100 mM DTT, 5% glycerol) to generate an input sample. 90% of the cells were resuspended in 0.2 mL lysis buffer containing 6 M Guanidinium-HCl, 100 mM K_2_HPO_4_, 20mM Tris-HCl (pH 8.0), 100 mM NaCl, 0.1% Triton X-100, and 10 mM Imidazole, and incubated on ice 20 min. Lysates were passed through a 30G needle five times. Purification of the His6-SUMO2 conjugates was performed on 50 μL of Ni^2+^-NTA agarose beads (Qiagen) prewashed with lysis buffer, and incubated for 2-3 h at room temperature with end-over-end rotation. The beads were washed once with 1 mL of lysis buffer, and three times with 1 mL of wash buffer containing 8 M urea, 0.1 M Na_2_HPO_4_/NaH_2_PO_4_ (pH 6.4), 0.01 M Tris-HCl (pH 6.4), 10 mM Imidazole, 10 mM β-mercaptoethanol, and 0.1% Triton X-100 before elution in 2X SDS loading buffer.

To detect conjugates exogenous ubiquitin conjugates, cells were co-transfected with plasmids (2.5 μg each) expressing His-Myc-ubiquitin (Li et al., 2010) and FH-AGO1 proteins or control plasmids and were treated with 10 μM MG132 for 7 h before harvesting. Ni^2+^-purified His6-ubiquitin (Ub) forms of FH-AGO1 followed by western blotting analysis were conducted as described above for the sumoylation assay.

To detect ubiquitin modifications on AGO1, cells were transfected with plasmids (2.5 μg each) expressing FH-AGO1 proteins or FH-EYFP (used as a negative control) and were treated with 10 μM MG132 for 7 h before harvesting. 24 h after transfection, cells were lysed in RIPA buffer (50 mM Tris-HCL, 150 mM NaCl, 1% Triton X-100, 1 mM EDTA, and complete protease inhibitor cocktail (Roche), pH 7.4) for immunoprecipitation. Cell lysates were incubated with Protein-G Sepharose beads coupled to the anti-HA antibody for 2-3 h at 4 °C. The beads were washed three times with RIPA and then processed for immunoblotting.

### Statistical analysis

Statistical tests were performed using two-tailed Student’s *t*-test to calculate P-values. P-value ≤ 0.05 was considered statistically significant. Results of statistical analysis are included in the corresponding figure legends.

### Ribosome fractionation

U2OS and RanBP2-dE3 cells were seeded at a confluency of 50-60% into 15 cm dishes and incubated for 24 h. The cells were then transfected with *IL6-li-HA* plasmid (15 μg) using GenJet-OS (SignaGen). 24 hours post-transfection, cells were treated with 100 μg/mL cycloheximide (Sigma) for 15 minutes, harvested using trypsin with 100 μg/mL cycloheximide, and washed two times in PBS with 100 μg/mL cycloheximide. Cells were then lysed in 1 mL of lysis buffer (20 mM HEPES-KOH (pH 7.4), 5 mM MgCl_2_, 50 mM KCl, 1% Triton X-100, 100 μg/ml cycloheximide, complete protease inhibitor cocktail (Roche), 1 μL DEPC) for 15 min. The lysates were cleared by centrifugation at 16,000 *g* for 10 min at 4°C and 400 μL of the lysates were layered over a 20-50% sucrose gradient in polysome buffer (20 mM HEPES-KOH (pH 7.4), 5 mM MgCl_2_, 125 mM KCl, 100 μg/mL cycloheximide). The solution was centrifuged at 36,000 rpm for 2 h in a SW41Ti rotor (Beckman Coulter) at 4°C. Gradients were fractionated and absorbance at 254 nm was measured using the Piston Gradient Fractionator (Biocomp) coupled with Bio-Rad EM1 UV monitor following the manufacture’s protocol.

RNA was isolated from each fraction using the Trizol reagent (Invitrogen) according to the manufacture’s protocol. Fractions representing free RNA, monosome, light polysomes (2-4 ribosomes), medium polysomes (5-7 ribosomes) and heavy polysomes (>8 ribosomes) were identified using the gradient profiles generated and were pooled together. *In vitro* transcribed Renilla *luciferase* mRNA (a final amount of 0.3 ng after pooling) was spiked into the fractions before RNA extraction as an external control. The RNA samples were treated with DNase I (Invitrogen), reversed transcribed into cDNA with SuperScript™ III Reverse Transcriptase (Invitrogen), analyzed by RT-qPCR (Applied Biosystems™ SYBR Green master mix) as per manufacture’s protocol. Primers used for RT-qPCR were, *α-tubulin* forward: CCAAGCTGGAGTTCTCTA; reverse: AATCAGAGTGCTCCAGGG; *IL6* forward: TGAGAGTAGTGAGGAACAAG; reverse: CGCAGAATGAGATGAGTTG; and Renilla *luciferase* forward: ATAACTGGTCCGCAGTGGTG; reverse: TAAGAAGAGGCCGCGTTACC. RT-qPCR data was analyzed using the ΔCT method where the relative expression of *IL6* and *tubulin* were normalized to that of Renilla *luciferase*.

### miRNA titration assay

U2OS and RanBP2-dE3 cells were seeded at a confluency of 20% in 6-well plates (0.3×106 cells/well) and incubated for 24 h. The cells were then transfected with 48, 24, 12, or 6 pmol of miR-644 Mimic (Dharmacon) or miRIDIAN miR-144 Mimic (Dharmacon) as a negative control using JetPRIME reagent (PolyPlus) according to the manufacturer’s instructions. Cell lysates were collected 48 h after transfection, proteins were separated by SDS-PAGE and immunoblotted for RanGAP1, GAPDH, and α-tubulin.

### Cell Fractionation

U2OS cells were seeded at a confluency of 50-60% into 15 cm dishes and incubated for 24 h. The cells were then transfected with *IL6-Δi-HA* plasmid (16 μg) using JetPrime reagent (PolyPlus). 24 hours post-transfection, cells were harvested using trypsin and washed two times in PBS. Cells were resuspended in Phi buffer (20mM HEPES-KOH (pH7.4), 150mM potassium acetate, 5mM magnesium acetate, complete protease inhibitor cocktail (Roche), 10mM PMSF and SUPERase In RNase inhibitor (Thermo Fisher)). The cells were lysed in Phi buffer with 0.5% Triton X-100 and 0.25% sodium deoxycholate for 1 min and spun at 1000g for 5 min at 4°C. The supernatant was kept as the cytoplasmic/ER fraction and was spun at 15,000g for 15 min to clear. Pelleted nuclei were washed two times to remove any unlysed cells.

### RNA-immunoprecipitation

Cell fractionation was performed as described above. The pelleted nuclei were lysed in high salt buffer (Phi buffer with 1% Triton X-100, and 125mM NaCl) for 30 min with rotation at 4°C. IP buffer (Phi buffer with 1 % Triton X-100) was then added to dilute out the salt. The fractions were then treated with DNase I (Thermo Fisher) at room temperature for 20 min and were spun at 10,000g for 10 min to clear. 5% of the cleared cytoplasmic/ER and nuclear fractions were collected and processed for immunoblotting. For the remaining, the fractions were split equally into two, mixed with 10 μg of control or anti-AGO1 synthetic antibodies (Na et al., 2016) and incubated overnight with rotation at 4°C. Flag M2 beads (Sigma) were blocked with 1% BSA, washed 3 times with IP buffer and incubated with the lysate-synthetic antibody mixture for 3 hours with rotation at 4°C. The beads were washed 5 times in IP buffer. 10% of the beads were collected and processed for immunoblotting. RNA was collected from the remaining beads with Trizol Reagent (Invitrogen) according to the manufacturer’s protocol and processed for RT-qPCR analysis as described above.

## ACKNOWLEDGEMENTS

We would like to thank J. Claycomb for valuable discussions concerning the work in the paper and for providing various reagents. We would like to thank L. Frappier, J. Vogel, T. Tuschl, and F. Zhang for plasmids. This work was funded by a CIHR grant to A. F. Palazzo (FRN 102725).

## AUTHOR CONTRIBUTIONS

All experiments were conceived and designed by Q. Shen, M. Truong, K. Mahadevan, Y.E. Wang, J. Wu and A.F. Palazzo. Reagents were prepared by H.W. Smith and C. Smibert. The experiments were performed by Q. Shen, M. Truong, K. Mahadevan, Y.E. Wang, and J. Wu. The manuscript was written by Q. Shen and A.F. Palazzo.

## Supplementary Information

**Figure S1. RanBP2 represses the expression of IL6 independently of splicing or the SSCR**

(A) Schematic of the *IL6* constructs tested. This includes an intronless version of *IL6* (*IL6-Δi*), a version containing the first endogenous *IL6* intron (*IL6-1i*) or the *ftz* intron (*IL6-1f*) both inserted at the endogenous first exon-exon boundary. (B-C) U2OS cells were infected with lentivirus that delivered shRNA1 against RanBP2 or control virus. Three days post-infection, cells were transfected with plasmids containing the indicated reporter genes. 18–24 h post-transfection cell lysates were collected and separated by SDS-PAGE. The level of each protein was analyzed by immunoblot for HA, and α-tubulin as a loading control (B). The levels of each HA-tagged protein and α-tubulin were quantified using densitometry analysis. The HA/tubulin ratio was normalized to *IL6-Δi* transfected control shRNA-treated cells and plotted (C). (D) Schematic of the *IL6* constructs tested. This includes a version of *IL6* where the endogenous SSCR was replaced with the mouse *MHC* SSCR derived from the *h2kb* gene (*MHC-IL6-Δi*). (E-F) As performed in (B-C) with each bar being the average of 3 independent experiments ± SEM. \**P* = 0.01–0.05 (Student’s *t*-test).

**Figure S2. Localization of RanBP2 variants**

WT U2OS, and RanBP2-dE3 cells were fixed and immunostained for RanBP2 and DAPI stained to visualize DNA. Note that the RanBP2-dE3 mutant localizes to the nuclear rim like the unmodified protein. Scale bar = 10 μm

**Figure S3. The SUMO E3-ligase domain of RanBP2 is required for IL6 translational suppression in HAP1 cells.**

The SUMO E3-ligase domain of RanBP2 was targeted by CRISPR/Cas9 in the haploid human HAP1 cells. (A) A schematic diagram of the region of the *RanBP2* gene targeted by CRISPR/Cas9 loaded with the guide RNA, “gRNA-dE3-1#”, whose sequence is shown in Figure 3B. Note that the insertion “ins” in the “RanBP2-E3 ins” cell line is indicated. (B) Sequencing of the single allele of the *RanBP2* gene in RanBP2-E3 ins cells. Note that the allele contained a 51 base pair insertion just downstream of the PAM sequence of the guide RNA. This insertion corresponds to a 17 amino acid insertion into the SUMO E3-ligase domain of RanBP2. (C-D) Wildtype (WT) HAP1 and RanBP2-E3 ins cells were transfected with plasmids containing an intron-containing version of *IL6-HA* and *H1B-GFP* and then cell lysates were collected 24 h post-transfection. IL6-HA was immunoprecipitated with mouse anti-HA antibody and protein G beads (Sigma), separated by SDS-PAGE and immunoblotted with a rabbit anti-HA antibody (top panel). For the detection of other proteins, cell lysates were directly separated by SDS-PAGE and immunoblotted with antibodies against GFP, RanBP2, RanGAP1 and α-tubulin. Note that RanBP2-E3 ins cells lacked sumoylated-RanGAP1 but expressed RanBP2 at similar levels to WT U2OS cells. IL6-HA and H1B-GFP protein levels were quantified using densitometry analysis and the ratio of IL6-HA/H1B-GFP was normalized to WT U2OS cells. Each bar is the average of 3 independent experiments ± SEM. \**P* = 0.01–0.05 (Student’s *t*-test).

**Figure S4. The RanBP2-responsive element is found in the 3′ end of the *IL6* 3′UTR**

(A) Schematic of the *IL6-1i*, *IL6-1i-3del1*, and *IL6-1i-3del2* constructs. 3del1 consists of the deletion of first 1-110 nucleotides of the *IL6* 3′UTR whereas 3del2 consists of the deletion of 111-439 nucleotides of the *IL6* 3′UTR. (B-C) Control and RanBP2 knockdown cells were transfected with HA-tagged *IL6-1i*, *IL6-1i-3del1*, and *IL6-1i-3del2* constructs. Cellular lysates were collected 24 h post-transfection, separated by SDS-PAGE and immunoprobed for IL6-HA and α-tubulin (B). The ratio of IL6-HA relative to tubulin was quantified using densitometry then normalized to control shRNA-treated cells transfected with *IL6-1i* (n = 4, mean ± SEM, \**P* = 0.01–0.05 NS indicates no significant (Student’s *t*-test)) (C).

**Figure S5. RanBP2 is required for elevated levels of AGO1 but not AGO2**

(A) U2OS cells treated with shRNA1 against RanBP2 were lysed, separated by SDS-PAGE, and immunoblotted with antibodies against AGO1, AGO2, RanBP2, RanGAP1, and α-tubulin. (B) WT U2OS, RanBP2-dE3 and RanBP2-dE3 cells which stably express GFP-RanBP2 with three ANE1 mutations were lysed, separated by SDS-PAGE, and immunoblotted with antibodies against AGO1, AGO2, RanBP2, RanGAP1, and α-tubulin.

**Figure S6. RanBP2 stabilizes overexpressed FH-AGO1**

(A-B) WT U2OS, and RanBP2-dE3 cells were co-transfected with *FH-AGO1* and *H1B-GFP.* 18 h post-transfection cells were treated with cycloheximide (CHX, 100 µM) in the presence of MG132 (10 µM) or DMSO for 7 hr. Cell lysates were collected, separated by SDS-PAGE, and immunoblotted with antibodies against HA, GFP, AGO2, RanBP2 and α-tubulin (A). FH-AGO1 and H1B-GFP protein levels were quantified using densitometry analysis and the ratio of FH-AGO1/H1B-GFP was normalized to DMSO-treated WT U2OS cells (B). Each bar is the average of 3 independent experiments ± SEM. *\*P* = 0.01–0.05, *\*\*P* = 0.001–0.01, n.s. indicates no significant difference (Student’s *t*-test).

**Table S1. ANE1-associated cytokines from case reports dating back to 1998**
| ANE1-associated cytokines | Number of 5'UTR Introns | Reference (Case reports that measured cytokine levels in patients) |
| --- | --- | --- |
| IL6 | None* | (Akiyoshi et al., 2006; Ichiyama, 2017; Ichiyama et al., 1998, 2003b, 2003a; Ito et al., 1999; Kawada et al., 2003; Kubo et al., 2006; Tabarki et al., 2013) |
| TNF-α | None | Ichiyama *et al*, 1998, Ichiyama *et al*, 2003a 2003b, Akiyoshi *et al*, 2006, Tabarki *et al*, 2013 |
| IL10 | None | Ichiyama *et al*, 2003a, Kubo *et al*, 2006, Kansagra *et al*, 2011 |
| IL15 | 2 | Kubo *et al*, 2006 |
| sTNFR1 | None** | Ito *et al*, 1999, Ichiyama *et al*, 2003a 2003b, Kubo *et al*, 2006 |
| IFNγ | None | Ichiyama *et al*, 2003a, Kubo *et al*, 2006 |
| IL1β | 1*** | Ichiyama *et al*, 1998, Ito *et al*, 1999, Kansagra *et al*, 2011 |
| IL2 | None | Kansagra *et al*, 2011 |
| IL5 | None**** | Kansagra *et al*, 2011 |

**Table S1. ANE1-associated cytokines from case reports dating back to 1998**

*- The major isoform lacks 5’UTR introns. There are three minor spliced isoforms, one the lacks a 5’UTR intron, one that uses an upstream transcriptional start site and whose extended 5’UTR has an intron, and a third that uses a downstream start codon and has a single 5’UTR intron.

**- The major isoform lacks 5’UTR introns. There are two spliced isoforms that contain the same 5’UTR but have been reported to use different start codons which are found in internal exons.

These extended 5’UTRs have two and five introns, respectively.

***- There is a reported minor spliced isoform that has a different 5’UTR which also contains one intron.

****- The major isoform lacks 5’UTR introns. There are three reported minor spliced isoforms, one that lacks 5’UTR introns, and the remaining two with a single 5’UTR intron.

