## Supplementary figures and images for "RanBP2/Nup358 enhances miRNA activity by sumoylating and stabilizing Argonaute 1"

### Figure S1

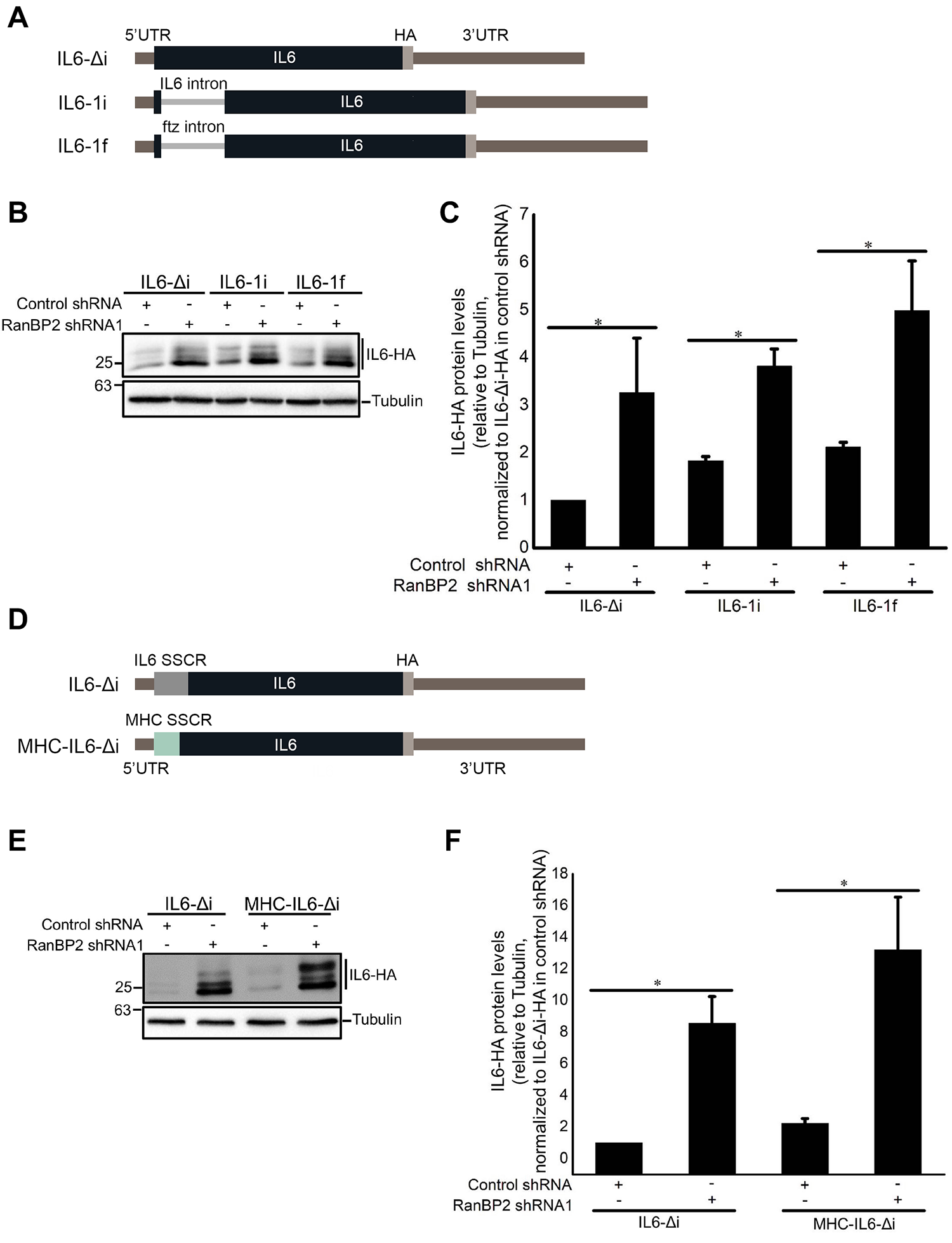

### Figure S2

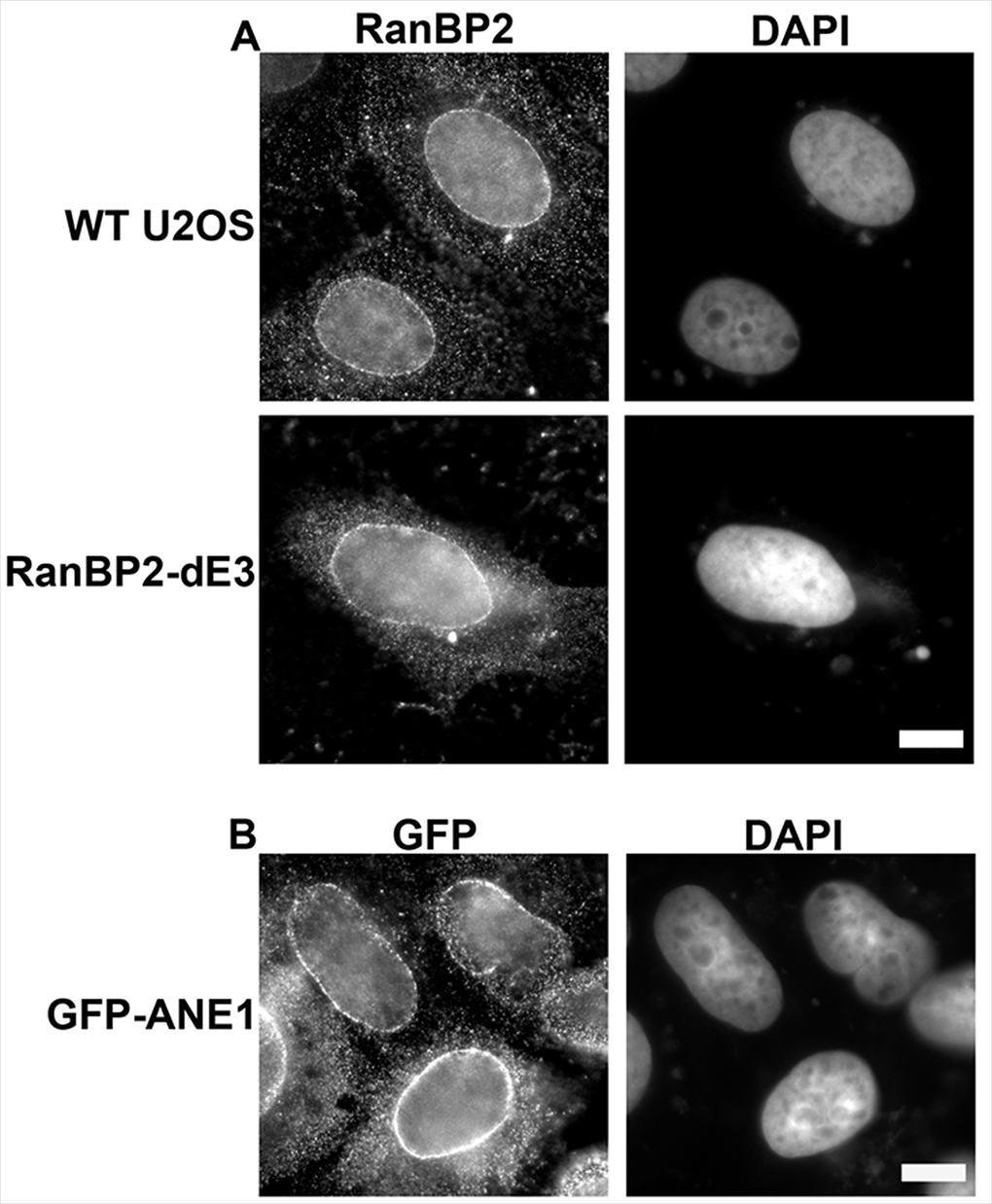

### Figure S3

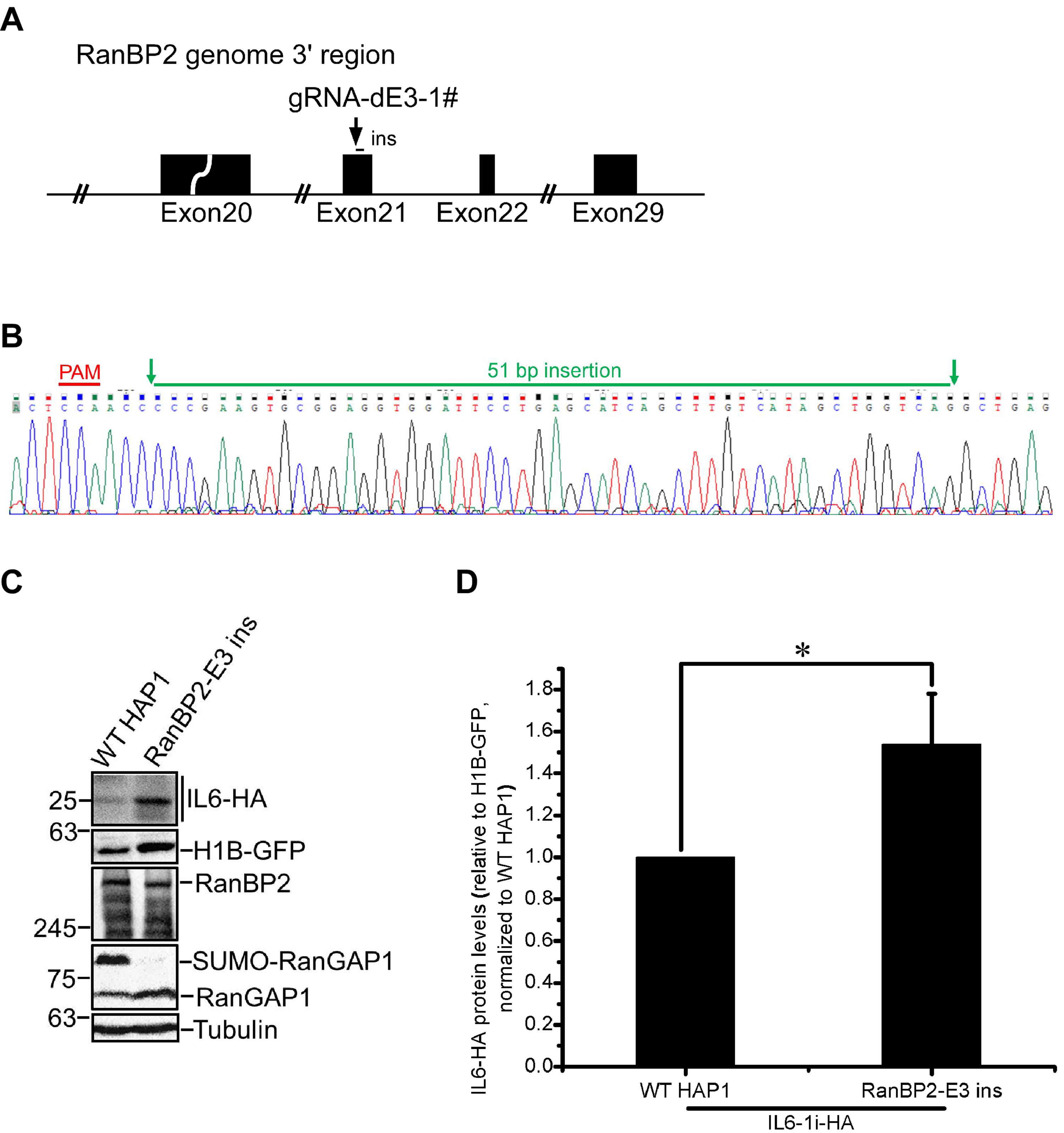

### Figure S4

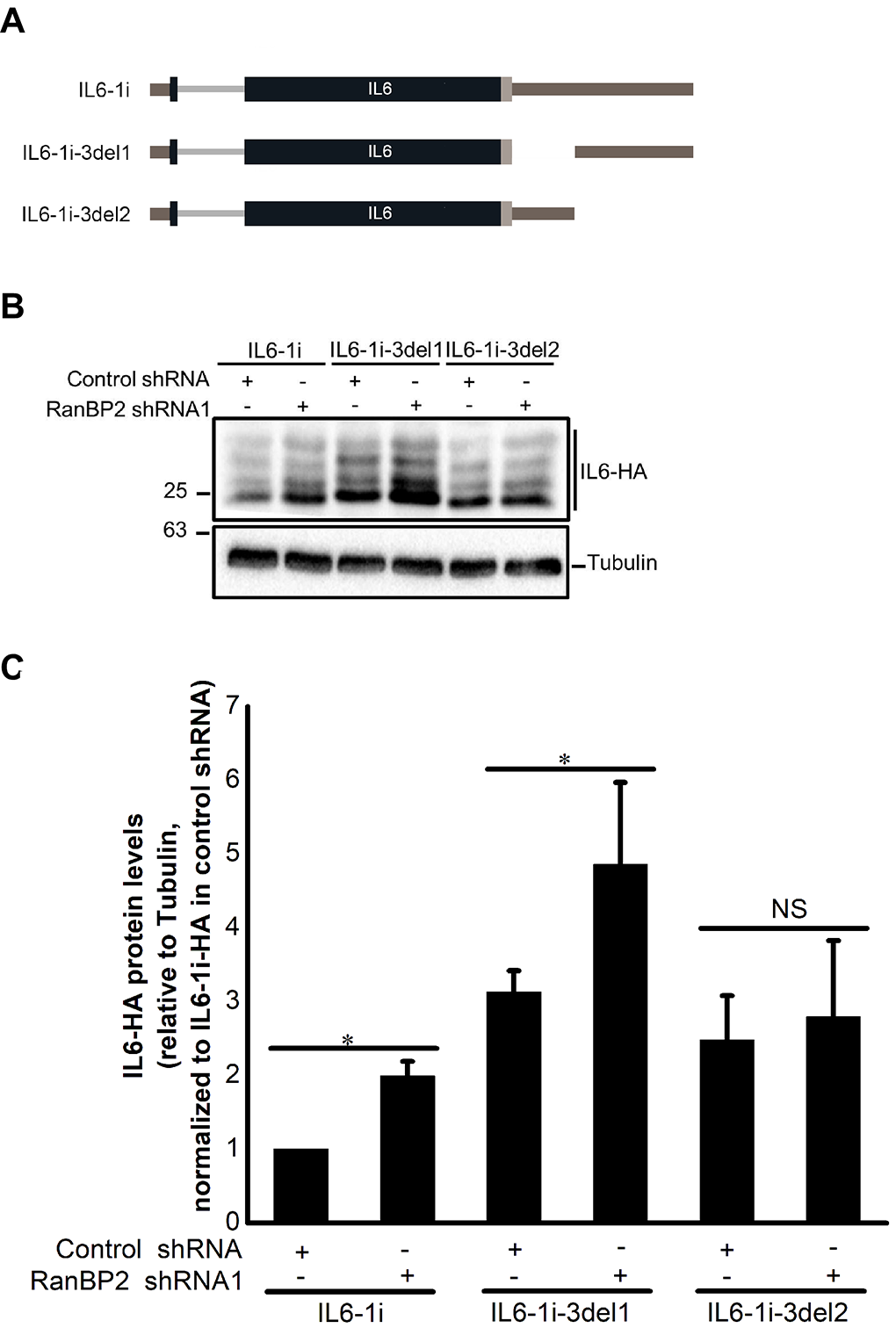

### Figure S5

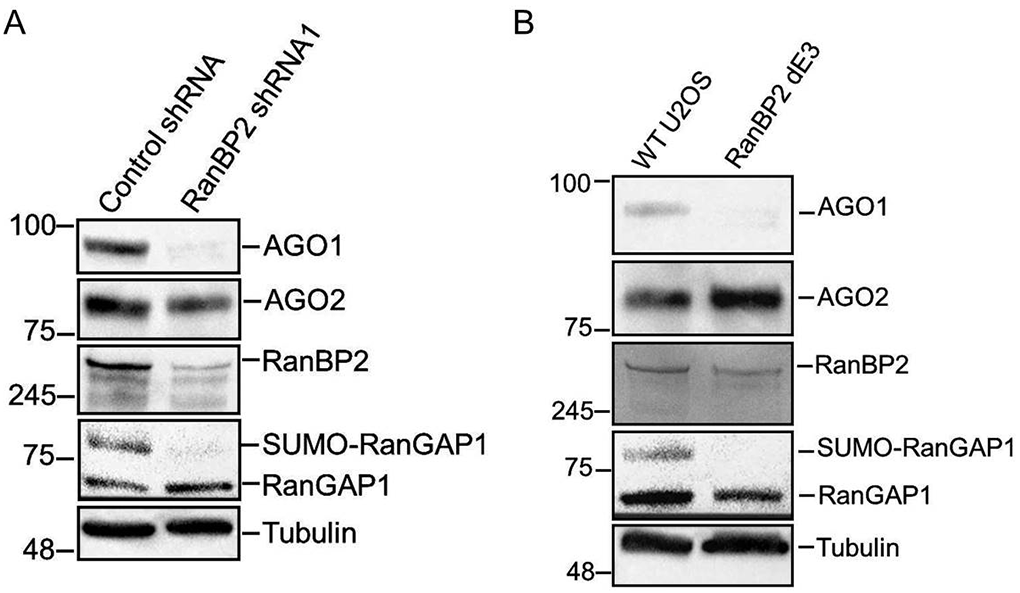

### Figure S6

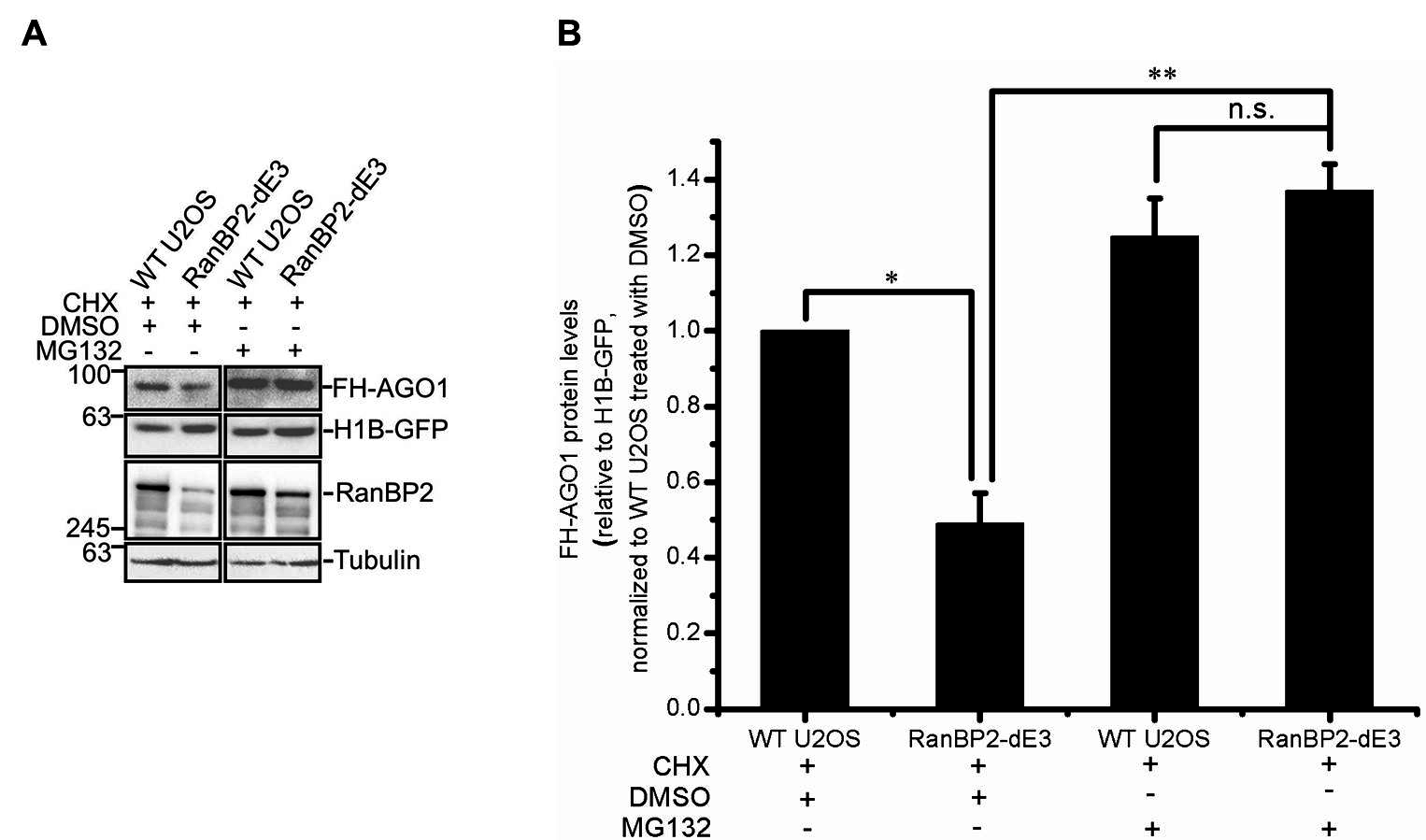
